# Feature selection and aggregation for antibiotic resistance GWAS in *Mycobacterium tuberculosis*: a comparative study

**DOI:** 10.1101/2022.03.16.484601

**Authors:** K.O. Reshetnikov, D.I. Bykova, K.V. Kuleshov, K. Chukreev, E.P. Guguchkin, V.G. Akimkin, A.D. Neverov, G.G. Fedonin

## Abstract

Drug resistance (DR) remains a global healthcare concern. In contrast to other human bacterial pathogens, acquiring mutations in the genome is the main mechanism of drug resistance for *Mycobacterium tuberculosis* (MTB). For some antibiotics resistance of a particular isolate can be predicted with high confidence knowing whether specific mutations occurred, but for some antibiotics our knowledge of resistance mechanism is moderate. Statistical machine learning (ML) methods are used in attempts to infer new genes implicated in drug resistance. These methods use large collections of isolates with known whole-genome sequences and resistance status for different drugs. However, high correlations between the presence or absence of resistance to drugs that are used together in one treatment regimen complicate inference of causal mutations by traditional ML. Recently, several new methods were suggested to deal with the problem of correlations of response variables in training data. In this study, we applied the following methods to tackle the confounding effect of resistance co-occurrence in a dataset of approximately 13 000 complete genomes of MTB with characterized resistance status for 13 drugs: logistic regression with different regularization penalty functions, a polynomial-time algorithm for best-subset selection problem (ABESS), and “Hungry, Hungry SNPos” (HHS) method. We compared these methods by the ability to select known causal mutations for the resistance to each particular drug and not to select mutations in genes that are known to be associated with resistance to other drugs. ABESS significantly outperformed the others selecting more relevant sets of mutations. We also showed that aggregation of rare mutations into features indicating changes of PFAM domains increased the quality of prediction and these features were majorly selected by ABESS.

**Impact statement:** Due to the high significance of the problem, many studies in the recent decade aimed to predict drug susceptibility/resistance of MTB from its genotype. Most of such methods were based on prior biological knowledge, e.g. consideration of mutations occurring in known genes involved in the metabolism of drugs. In our study, we estimated to what extent ML methods could extract de novo biologically relevant associations of mutations with resistance/susceptibility to drugs from large datasets of clinical MTB isolates. As a criterion of accuracy we used the known experimentally verified associations of mutations in MTB genes to corresponding drugs. The most accurate approach from the benchmarked ones addressed the most of these known genes to proper drugs. The result of feature selection was robust despite the presence of population structure with strong phylogenetic and geographic signals in the dataset. Also, we designed an original approach for aggregation of rare mutations and demonstrated that it improved classification accuracies of ML models. To our knowledge, this study is the first comparison of modern feature selection methods applied to genome-wide association studies (GWAS) of MTB drug resistance.

**Data Summary:** The dataset unifies characterized whole-genome sequences of *M. tuberculosis* from multiple studies [1–10]. Short Illumina reads are available in public repositories (SRA or ENA). Sample ids, phenotypes and links to the source papers are summarized and listed in Table S1. The dataset and the source code can be downloaded from the GitHub repository: https://github.com/Reshetnikoff/m.tuberculosis-research-code

## Introduction

Tuberculosis (TB), caused by *Mycobacterium tuberculosis* (MTB), remains the leading cause of death from single infectious agents in spite of decreasing annual numbers of TB deaths in recent years [11]. TB is a curable disease, which, in most cases, could be effectively treated by a complex of drugs, there is a great concern about increasing fraction of isolates resistant to the most potent anti-TB agents rifampicin (RIF) and isoniazid (INH) called multidrug resistant (MDR) as well as extensively resistant (XDR) isolates which are additionally resistant to any aminoglycosides and fluoroquinolones [11]. According to the WHO estimates, the numbers of multidrug-resistance tuberculosis (MDR-TB) and rifampicin resistance tuberculosis (RR-TB) cases detected had increased from 2018 to 2019 by 10% [11]. The average efficiency of treatment of MDR-TB over the world is about 60% and for XDR-TB is about 20%, and the prognosis of the life span for treatment failures is negative [11]. Today, the two basic schemas of TB therapy are applied [12], the naive patients are effectively treated by a combination of first-line drugs rifampicin, isoniazid, pyrazinamide (PZA), ethambutol (EMB), and streptomycin (STR), and relapse cases are treated by the second-line drugs fluoroquinolones: ofloxacin (OFX), levofloxacin (LEV), moxifloxacin (MOX) and ciprofloxacin (CIP) and injectable drugs kanamycin (KAN), amikacin (AMK) and capreomycin (CAP). KAN and CAP are no longer recommended for TB treatment [13]. There are other less effective second-line anti-TB drugs: ethionamide (ETH)/prothionamide (PTH), cycloserine (CS), P-aminosalicylic acid (PAS).

Early detection of resistant cases is necessary to correct therapy and prevent transmission of resistant strains because patients infected with drug resistant strains are harder to treat and sometimes easier to transmit disease [14]. Acquisition of mutations in genes employed in processes which are disturbed by drugs is a main mechanism of drug resistance in MTB [15]. Moreover, in some cases if MTB is resistant to one drug, it acquires resistance to other drugs easier [16].

Measuring the minimal concentrations of a substance inhibiting bacterial growth is the ‘gold standard’ for assessing resistance/susceptibility levels for drugs [17]. Because MTB is a slow-growing bacteria, phenotypic tests are time-consuming, expensive, and could be performed in laboratories with high biosafety levels only. Thus, the genotypic tests have been gradually replacing it. PCR-based molecular tests could be used for MTB DNA extracted directly from clinical samples. MTB GeneXpert MTB/RIF, line probe assays MTBDRplus and MTBDRsl, detect the presence of frequent mutations in a few drug associated genes within two hours and have high DR prediction accuracy for RIF and INH, but lower for other drugs [18–20]. Meanwhile, molecular methods based on sequencing allow for detection of rare mutations and have higher accuracy [21]. Despite the fact that sequencing tests usually require the MTB culturing stage, they are less time and cost consuming than standard laboratory diagnostics [22]. In Great Britain [23] and in some other high-income countries [24] WGS-based tests are adopted for use for drug susceptibility testing before prescription of therapy. WHO recently released the technical guide for use of Next Generation Sequencing (NGS) for the detection of mutations associated with MTB drug resistance [13, 25, 26].

Most methods of MTB phenotype prediction by genotype utilize the direct association (DA) approach based on the interpretation of mutations as benign or causal [21]. Prior knowledge of genes, which products are likely implicated in drug action, allows the identification of drug-associated causal mutations. A pioneering DA approach was developed by Walker et. al. [9]: for each drug the method iteratively identifies mutations in a relevant subset of preselected genes which explain resistant phenotype of at least one previously unexplained isolate in the training set. Some other catalogs of resistance causal mutations were developed based on thorough literature reviews [24, 27, 28], machine learning methods [29] and comparative genomics [30]. Recently WHO made standardized methodology to compile a catalog of mutations in the preselected MTB genes best explaining drug phenotypes [13].

Direct association models perform well for some drugs, e.g. rifampicin and isoniazid, which action mechanisms are well known, however, for some drugs, e.g. pyrazinamide, ethambutol, fluoroquinolones, mutations in known genes do not always explain phenotypes well [21, 31].

Many approaches were used to identify new loci in the MTB genome associated with drug resistance by genome-wide association studies (GWAS) that process datasets of almost complete-genome genotypes with characterized drug phenotypes. Some methods are based on the detection of sites in the genome with recurrent mutations in phylogenetically unrelated strains [1–3, 32, 33], others use machine learning (ML) approaches: k nearest neighbors [34], regularized logistic regression [34–36], support vector machine [4, 35, 37], gradient-boosted trees [34, 38], random forest [4, 34], mixture models [4] and linear mixed models that account for the clonal structure of MTB population [3, 5, 39]. New methods are constantly being invented, for example, the one based on boolean compressed sensing [40]

Although a few loci identified in GWAS studies were experimentally confirmed as causal [2, 32, 33] others require verification and may be spuriously associated. Likely, the number of potential DR causal loci is large [5], e. g. in experiments with *E. coli*, high levels of resistance could be achieved by modulation of expression of a large number of functionally diverse genes [41], therefore, discovering numerous causal mutations is a hard objective.

Compared to the DA approach, the accuracy of drug phenotype prediction could be significantly improved by the application of ML for mutations occurring in known drug-associated genes [6, 21, 29, 37, 42, 43]. The improvement is partly achieved due to high correlation of DR phenotypes for drugs usually prescribed in common [6, 42], so identification in an isolate a specific mutation that causes resistance to one drug informs model about high probability of co-resistance to other drugs, e.g. the mutation *katG* S315T conferring resistance to INH also usually arises before acquisition of RIF resistance [44, 45] and is a good predictor for MDR [46, 47].

The model trained on data with certain proportions of MDR/XDR could be less accurate in predicting resistance for the data with different proportions of MDR/XDR [6] which may lead to the poor prediction quality of the models trained on clinical isolates for in-vitro selection experiments or for regions with different proportions of resistant TB. Therefore, such models will have high prediction accuracies only if regimens of usage of drugs will be the same in the future. But the WHO recommendations for treatment are constantly changing and so does the level of drug resistance in the population. Also, models based on resistance cooccurrence are unable to capture rare causal mutations because covariates would mask weak signals [42]. Finally, such models hardly produce new knowledge about the mechanisms of drug resistance.

Along with the resistance co-occurrence, the curse of dimensionality is one of the major problems in GWAS. Yang et al. among the others [6, 34] applied principal component analysis (PCA) to MTB DR drug prediction reducing dimensions, but separability between INH-resistant and susceptible classes was poor [42]. Artificial neural networks, including deep learning and autoencoders, are widely treated as automatic feature generation and were successfully applied to MTB DR prediction [21, 43]. These models, potentially, account for complex interactions between mutations. However, the improvement of prediction accuracy of such complex models in comparison to linear regression models is modest or insignificant [21] while interpretation of such models is nontrivial. Matrix factorization methods are another example of feature generation for dimensionality reduction. This approach could be unified with feature selection, as it was done recently in [48] for MTB DR prediction.

Aggregation of mutations with the subsequent exclusion of rare mutations is another way of dimensionality reduction by feature generation. It is typically done by constructing features indicating the presence of mutations in a particular gene or in some regulatory region [21]. Such indicators reduce dimensions without loss of interpretability and allow accounting for mutations that were not present in the training set. In this study, we tested HMM models from the PFAM database as a mechanism of aggregation.

Feature selection techniques try to select a subset of initial features, optimal in some sense. For large dimensions, however, enumeration of subsets is computationally intractable. Different methods vary in heuristics they use to evade the full enumeration. In this paper, we concentrate on feature selection with linear models. We compare multiple selection methods both in terms of prediction accuracy and biological relevance. We have chosen logistic regression with L1, minimax concave penalty (MCP) and smoothly clipped absolute deviation (SCAD) regularization, elastic net, ‘Hungry, Hungry SNPos’ (HHS), and a polynomial-time algorithm for best-subset selection problem (ABESS). A brief description of these methods is provided in the “Overview of the feature selection methods” (see Supplementary Materials).

In this study we benchmarked multiple feature selection methods mentioned above on a large dataset and compared their ability to choose biologically relevant genes and associate them with the proper drugs while being trained on the samples demonstrating resistance co-occurrence. We showed that the most popular feature selection techniques tend to assign genes which are known to be associated with one drug to the other drugs. We also proposed a two step feature selection procedure based on ABESS, the winner of our benchmark, which can find new associations.

We also tried to fight with the curse of dimensionality by constructing aggregation features based on PFAM domain annotations of proteins and demonstrated the usefulness of these features. Figure S1 summarizes the main steps of our study.

Geographic heterogeneity of the dataset is another important factor in GWAS. The dominant resistance mechanism can vary in different regions and it was shown recently that ML models trained on homogeneous datasets perform badly on the other homogeneous datasets [60]. Thus, training on heterogeneous datasets is more robust, but geographic or lineage markers can be associated with the resistance even in heterogeneous datasets.

Ignoring the population structure can lead to finding of the spurious associations especially in cases when there are benign mutations which are highly correlated with these markers. We tested our dataset on the presence of population structure and showed that there is a strong geographic signal, but our selection strategy is robust to it.

## Methods

### Dataset preparation

The dataset unifies phenotypically characterized whole-genome sequences of *M. tuberculosis* from multiple studies [1–10]. Illumina reads were trimmed and mapped onto the H37Rv reference genome. Variants were called with GATK HaplotypeCaller and multiple filtering steps were applied.

Nucleotide sequences of each protein-coding gene were translated and aligned to the reference protein sequences with extreme care to take into account possible frameshifts, start or stop codon loss, etc. The mutated gene was considered nonfunctional if it was 50% shorter or 30% longer than the reference protein: no mutations were considered in such genes. Finally, we obtained a list of mutations relative to the reference genome: mutations in noncoding regions, missense mutations, loss of function mutations, and aggregated features (see below). Synonymous mutations were excluded from the analysis.

The final dataset was split into smaller sets containing only samples with known phenotypes for each antibiotic. Both aggregation and machine learning were performed separately for each dataset. For each drug the dataset was randomly split into five non overlapping subsets (folds). Each fold was used as the testing set one at a time, while the union of the others served as the training set. Below we call every such partitioning “the dataset split”. For every dataset split the testing fold is called “the testing partition” and the union of the others is called “the training partition”

See “Dataset and raw data processing” in Supplementary Methods for details.

### Aggregation of mutations

We examined three types of aggregated features: an indicator of the presence of any mutation in a gene, an indicator that the gene is broken, i.e. it has a loss of function mutation, and features indicating alternation of PFAM domains.

If there is any mutation in a gene then the indicator of the presence of mutations in this gene takes the value of “1” and “0” otherwise. Further, this approach is called “gene aggregation”. “Broken gene” is another binary feature. Gene is considered to be ‘broken’ if the length of its open reading frame (ORF) is decreased due to mutations by half or increased by 30% relative to the annotated ORF length.

To generate PFAM domain features we used pre-trained models available for 3369 *M. tuberculosis* genes in the PFAM database. Using these models for each genome sequence we computed domain scores which were probabilities that domain sequences had been generated by corresponding domain HMM models. We then converted scores into binary variables selecting for each domain and for each drug separately a threshold optimizing separation of resistant and susceptible sequences in the training partition of each dataset split. If a domain score is over the threshold then the domain is considered to be changed and the corresponding binary variable takes the value of “1” and it takes the value “0” otherwise (see “PFAM domains based feature generation” in Supplementary methods).

### Estimating the impact of aggregation

To measure the effect of aggregation we evaluated results of classification of genotypes into resistant and susceptible phenotypic groups by logistic regression with L1 regularization using different feature set combinations. The basic feature set comprised SNPs and Indels without aggregation. Three other sets contain aggregating features and mutations that happened more often than 2 times: 1) SNPs and Indels with PFAM features and excluding rare mutations; 2) SNPs and Indels with “broken gene” features and excluding rare mutations; 3) SNPs and Indels with “gene aggregation” features and excluding rare mutations. In these three sets and everywhere below filtering of rare mutations was applied after the generation of aggregated features.

ROC AUC and F1 scores were computed by 5-fold cross-validation: the training partitions of dataset splits were used for training of logistic regression, the scores computed on testing partitions were averaged. Statistical significance of the differences of scores between the models trained on different feature sets was estimated by repeating this procedure 100 times and calculating Wilcoxon paired rank test between two vectors of length 100 of corresponding scores. The null model assumes that there is no difference between the performance of logistic regression on different feature sets and the difference of scores obtained on 5-fold cross-validation is explained only by the lucky dataset split. We considered the differences to be significant if the p-value with Bonferroni correction to the number of drugs was lower than 5%.

The schema of the comparison of feature aggregation strategies is presented in Fig. S2.

### Feature selection and model quality evaluation

The performances of methods: of regularized logistic regression (L1, MCP and SCAD), elastic net, HHS and ABESS, were estimated by 5-fold cross-validation. Again, the dataset was divided into five subsets referred further as folds. Union of four folds was used as a training set, while the fifth fold was used as a testing set. Each fold was used once as a testing set, as a result, different lists of mutations were selected for five splits of the dataset into training and testing sets.

After the training, a list of features that have non-zero coefficients as a result of the training of each method on each training subset was obtained. For each feature, the number of times that this feature has been selected by each particular method was computed.

We defined a feature to be majorly selected if it was selected in three or more dataset splits. For each such feature Fisher exact test p-value was computed on the testing part of the first dataset split. Ordinary logistic regression was trained using these features and ROC AUC, sensitivity, specificity, NPV and PPV were computed.

To search for additional features associated with the drug phenotypes that were missed by the described procedure of application of ABESS for the five-fold dataset splits we performed the second iteration of ABESS to the subset of data unexplained at the first iteration. For the second iteration of ABESS, we removed from the dataset resistant isolates correctly classified on the previous iteration and some randomly selected susceptible isolates to maintain the class balance. All features which were majorly selected for at least one drug with positive coefficients at the first iteration were completely removed, enforcing the method to search for the new associations. After the training of ABESS, the feature selection process was repeated. The same procedure was performed for HHS.

See Supplementary Methods for details.

### Accounting for the population structure

It is adopted widely that due to clonal evolution of bacteria some multicomponent genetic features associated with resistance could be shared by the groups of strains having common ancestry. Such groups will comprise distinct clades on the phylogenetic tree, which may be associated with geographic regions, may contain genetic lineages or be part of larger lineages. Therefore genetic markers of such clades could be spuriously associated with the resistance by any statistical learning method. To investigate if there are any subsets of phylogenetically related samples in our dataset such that the shared ancestry with them is a good marker of resistance we have built a maximum likelihood phylogenetic tree for all isolates included in our study (see Supplementary Methods for details). Separately for each drug we pruned all isolates that did not contain the phenotype information and obtained the corresponding subtree. We then applied TreeBreaker [61] to each subtree to infer the clades having the prevalence of resistant samples significantly different from the parental clades. The branches of each subtree having posterior probabilities of prevalence switches larger than 0.5 were used as structural binary features each of which took values equalled to one for all descendent isolates and to the zero for all other isolates.

Geographic locations of isolates were obtained from NCBI BioSample records. If such data was missed in a BioSample record the location of an isolate was assigned based on text descriptions in the corresponding BioProject record if possible. We used the information about a country where an isolate had been sampled for further analysis. To test if a distribution of isolates belonging to each clade identified by the TreeBreaker by country categories was significantly different from the distribution by categories of all isolates brought together we computed the sum of squares of differences of isolate frequencies within the country categories belonged to the clade and frequencies of isolates in the entire tree. We computed p-values for differences of distributions of isolates by country categories for the clades identified by the TreeBraker by multiple permutations of location labels of isolates. Bonferroni correction for the number of applied tests for all subtrees together was used for the p-values. We considered significant p-values below the Bonferroni corrected 5% threshold.

Isolates were assigned to the MTB genetic lineages based on the catalog of the TB-Profiler lineage-defining mutations [62]. To test if the distribution of isolates belonging to each clade identified by the TreeBreaker by lineages was significantly different from the distribution by lineages of all isolates brought together we applied the similar procedure as for the geographic location test.

Isolates that had no location or lineage information were ignored in the corresponding tests.

See Supplementary Methods for details.

## Results

### The most of mutations are rare

Short reads and phenotypes for 13 drugs were obtained from publicly available sources (TableS1) [1–10]. After subsequent data preprocessing our dataset comprised 12333 wholegenome sequences for which binary phenotype (resistance or susceptibility) for at least one of 13 antibiotics was known.

For all antibiotics, the number of unique mutations significantly exceeds the number of samples. Moreover, most of the mutations are rare: they appear in the dataset less than for three isolates. Such mutations comprise more than 67% of all unique mutations for all drugs (Table 1).

**Table 1.**
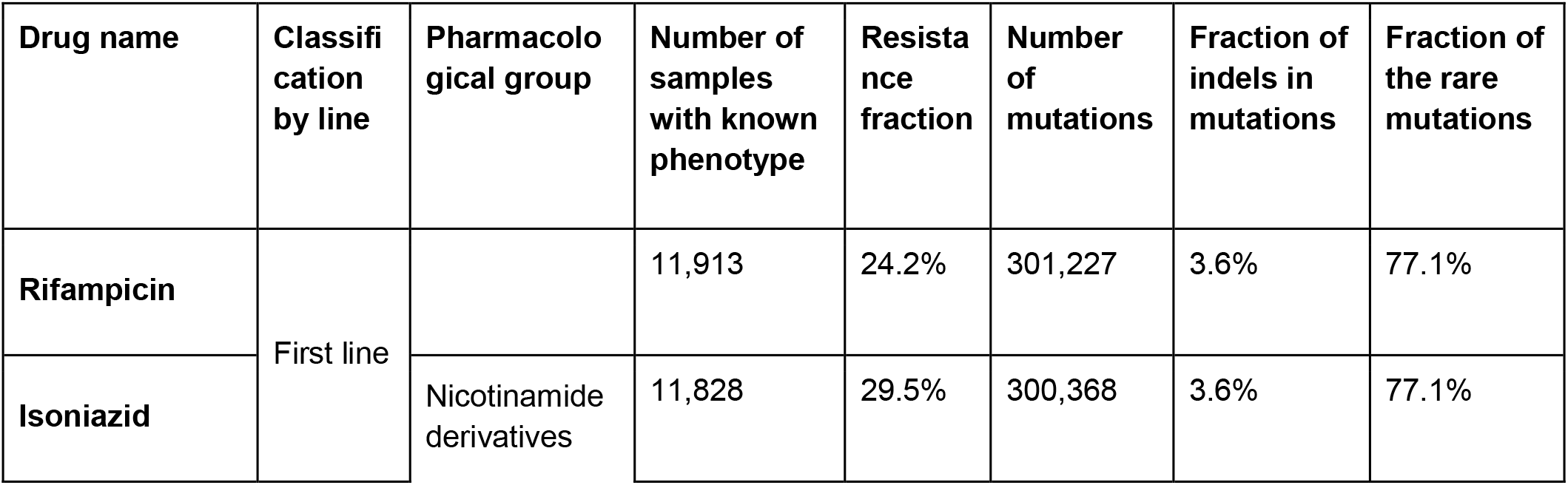
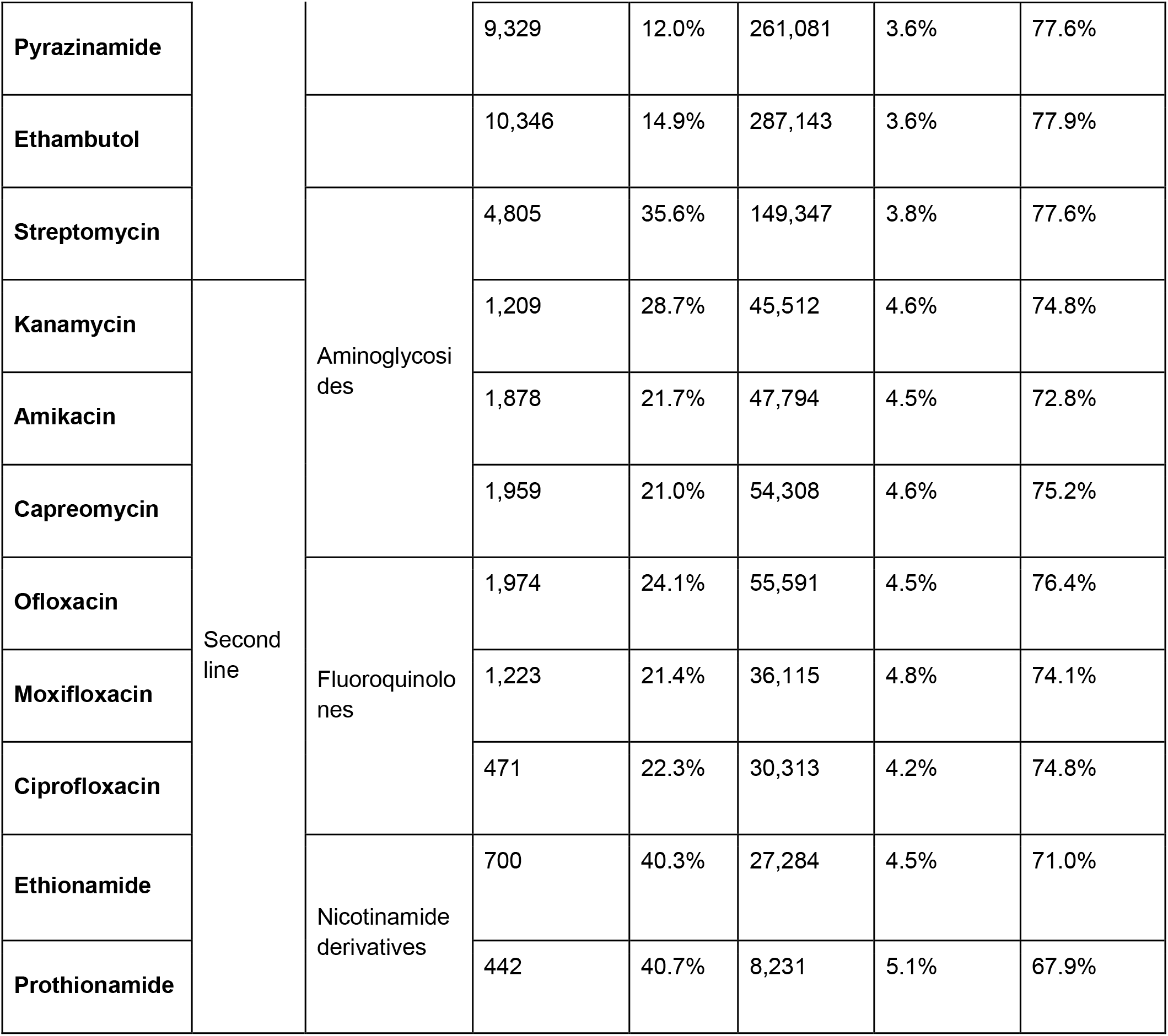
The descriptive statistics of the dataset. We call a mutation rare if it occurs less than 3 times in the dataset.

### Most of the drugs demonstrate coresistance patterns

To illustrate and quantitatively characterize the coresistance we have computed Pearson’s correlations for all pairs of drugs. The correlation was computed on the samples phenotypically characterized for both drugs in a pair. The resulting heatmap is presented in Fig. 1. Many pairs of phenotypes turn out to be strongly correlated (Fig. 1). High correlations are expected for drugs from one pharmacological group such as aminoglycosides (Amikacin, Kanamycin, Streptomycin, Capreomycin) and fluoroquinolones (Moxifloxacin, Ofloxacin, and Ciprofloxacin). Aminoglycosides share three known genes associated with resistance (*rrs*, *tlyA*, *gidB*) and fluoroquinolones share two (*gyrA*, *gyrB*) (Table 2). Another cause of high correlation is joint antibiotic treatment, which is, probably, the case for Rifampicin and Isoniazid, as well as for all pairs of the first-line drugs.

**Fig 1.**
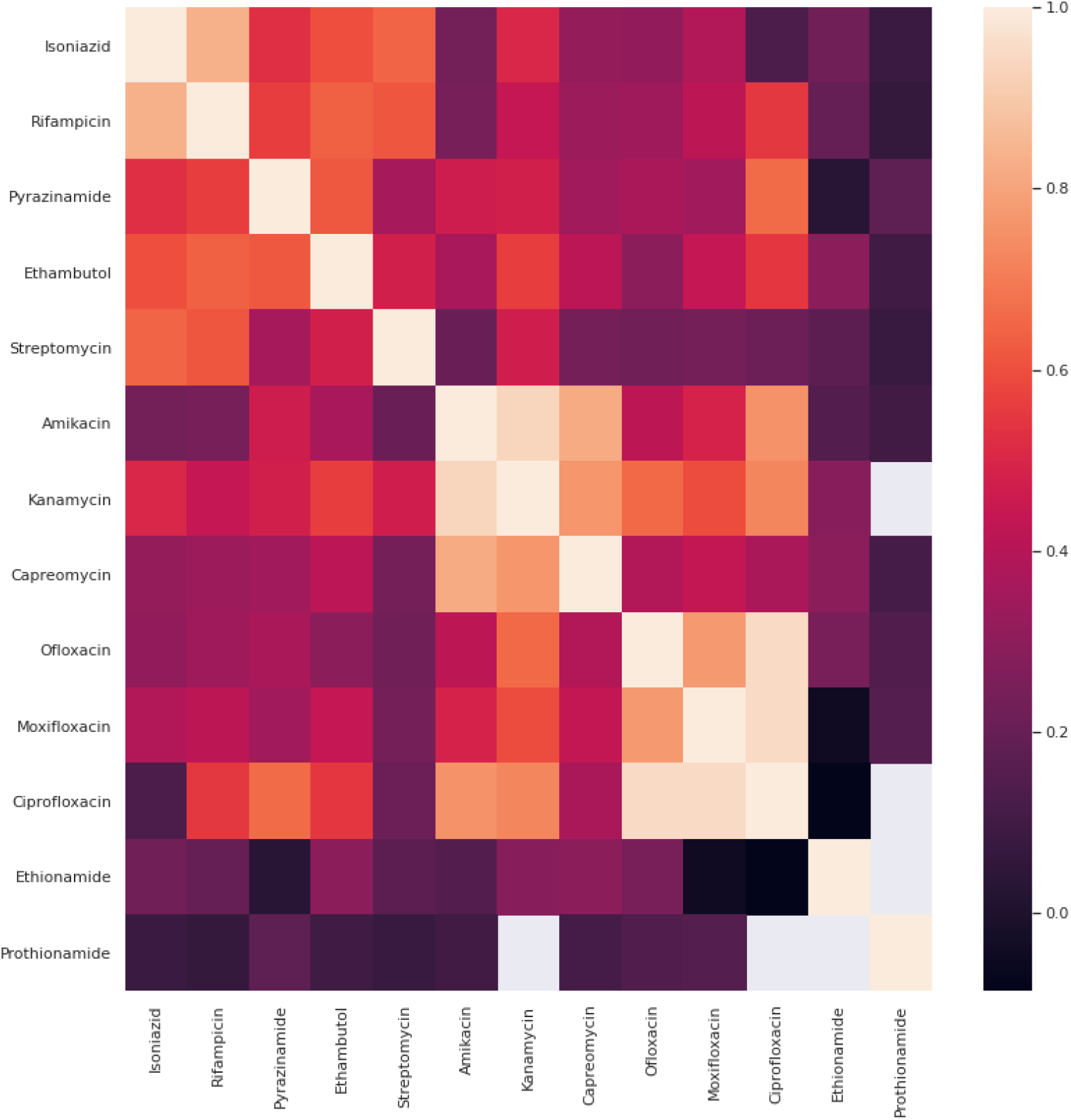
The heatmap of Pearson correlation between each drug phenotype pair. For each pair only isolates with both phenotypes being known were used.

**Table 2.**
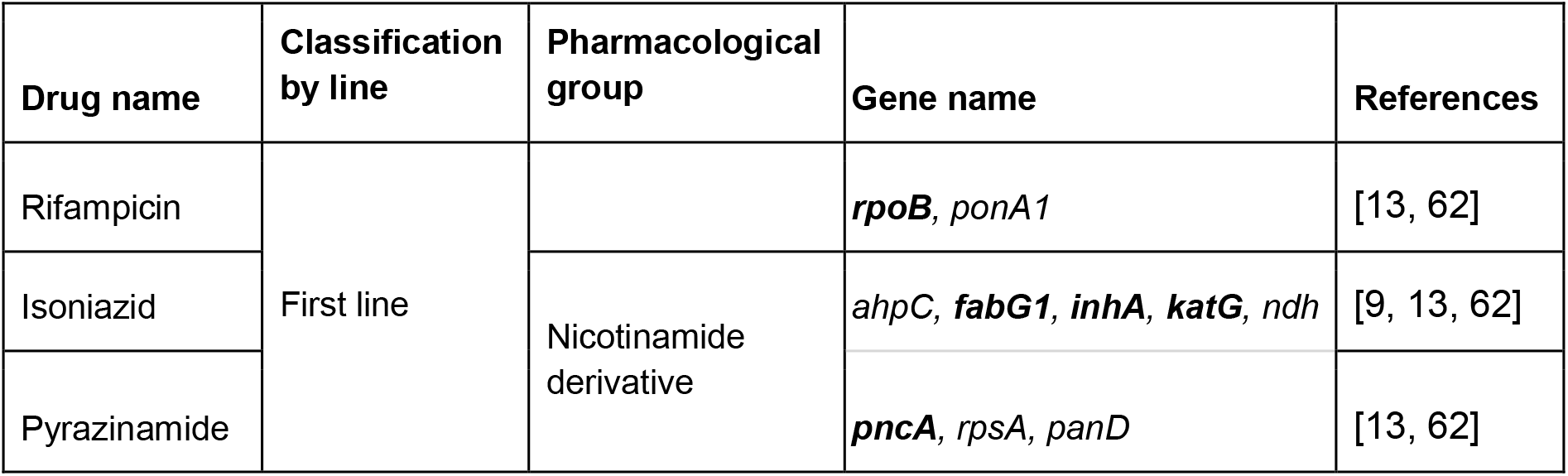
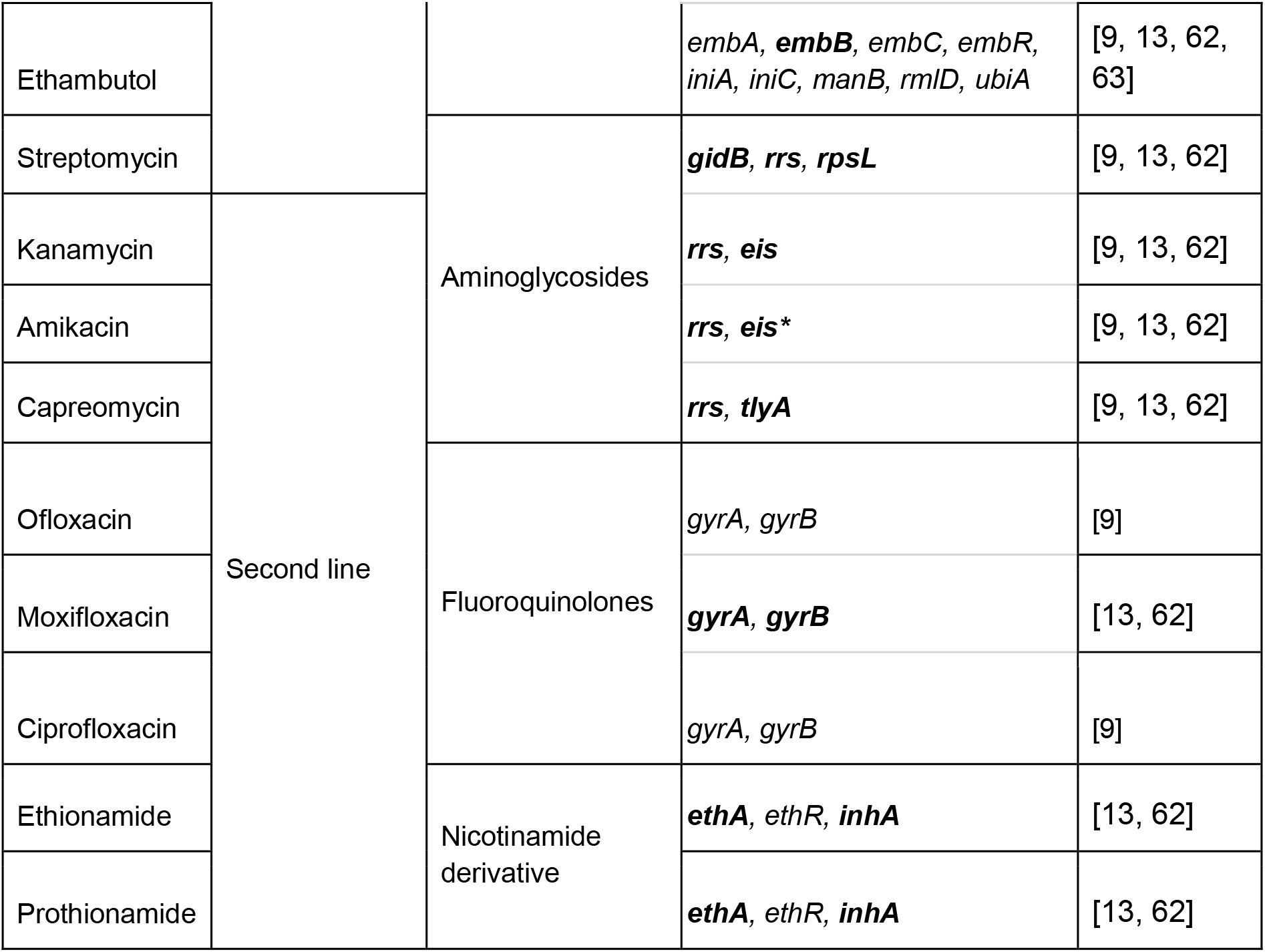
Genes associated with resistance to drugs used for the tuberculosis treatment according to previous studies. Genes from the 2021 WHO catalog are marked in bold. * marks eis gene which is present in WHO catalog but not in the other sources.

### Aggregation of mutations improves the prediction quality

To measure the effect of aggregation we evaluated results of classification of genotypes into resistant and susceptible phenotypic groups by logistic regression with L1 regularization using different feature set combinations (see “Aggregation of mutations” and “Estimating the impact of aggregation” in “Methods”). Four feature sets were involved: SNPs and Indels; SNPs and Indels with PFAM features and excluding rare mutations; SNPs and Indels with “broken gene” features and excluding rare mutations; SNPs and Indels with “gene aggregation” features and excluding rare mutations. Table S2-S7 presents the results of pairwise ROC AUC and F1 scores comparison of sets with aggregation to the basic set comprising SNPs and Indels. We performed comparisons for each drug separately and for drug groups (first-line, second-line, aminoglycosides, fluoroquinolones and all drugs together). For groups, the values of scores are arithmetic means of the corresponding scores obtained for drugs within groups. Sets with aggregated features demonstrated higher prediction quality for some drugs and the difference was usually statistically significant (Fig. 2, Table S2-S7). The differences of values of ROC AUC are moderate, but, frequently, have large statistical significance due to small variances for all feature sets (Table S8). The feature sets including the PFAM domain features have higher AUC scores for all drug groups except fluoroquinolones and for six drugs individually (Table S2). “Broken genes” feature wins in pairwise comparisons for 9 from 13 drugs (Table S3) and “gene aggregation” - only for 5 from 13, while for 7 drugs the result is the opposite (Table S4). The results for F1 score are consistent with ROC AUC in general (Table S5-S7). We compared PFAM domain features with gene aggregation: PFAM domains-based aggregation outperforms gene aggregation for most drugs both in terms of ROC AUC and F1 score (Table S9-S10). We kept PFAM domain features and “broken genes” features in all further experiments.

**Fig 2.**
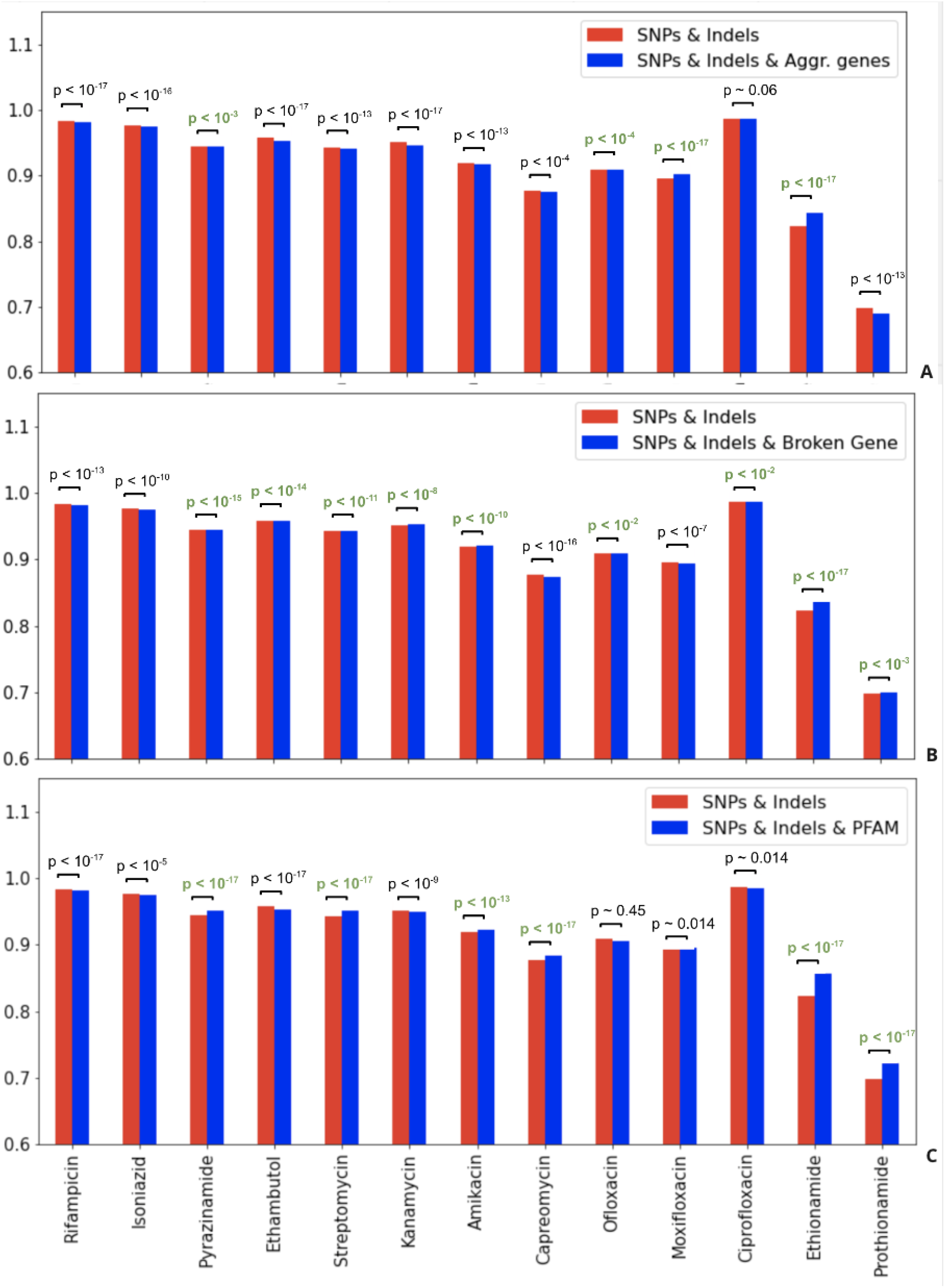
The comparison of mean ROC AUC scores of logistic regression evaluated on different combinations of features sets for drug groups between: A. “SNPs & Indels’’ and “SNPs & Indels & PFAM domains excluding rare mutations”; B. “SNPs & Indels” and “SNPs & Indels & “broken genes” excluding rare mutations”; C. “SNPs & Indels” and “SNPs & Indels & “gene aggregation” excluding rare mutations”. Wilcoxon signed-rank test was used for evaluating statistical significance. P-values corresponding to drug groups that demonstrated statistically significant increase in AUC due to aggregation are colored in green.

### Comparison of the feature selection algorithms

In this study, we explored six methods for the selection of causal DR mutations: logistic regression with L1, MCP, and SCAD regularization, elastic net, HHS, and ABESS. To test the performances of methods the datasets were split into 5-fold for cross-validation (see Methods). The list of mutations selected in at least one split by each method is listed in Table S11-S16 along with the numbers of splits in which it was chosen.

ROC AUC was computed to compare prediction quality of feature sets selected by different methods and the DA method based on WHO catalog [13]. Selected features obtained by the logistic regression with L1, SCAD, MCP and elastic net regularization techniques showed predictions of similar quality, whereas HHS was the worst one except for Capreomycin and Amikacin. The DA method based on WHO catalog for all drugs showed the lower quality of predictions compared to all methods except for the HHS which results were the worst for the most drugs (Fig. 3, Table S17-S23). This was expected since the catalog was constructed by applying very stringent statistical thresholds to label mutations as “associated with resistance” leading to higher specificity and lower sensitivity (Table S17-S23) and thus being too conservative in its predictions.

**Fig 3.**
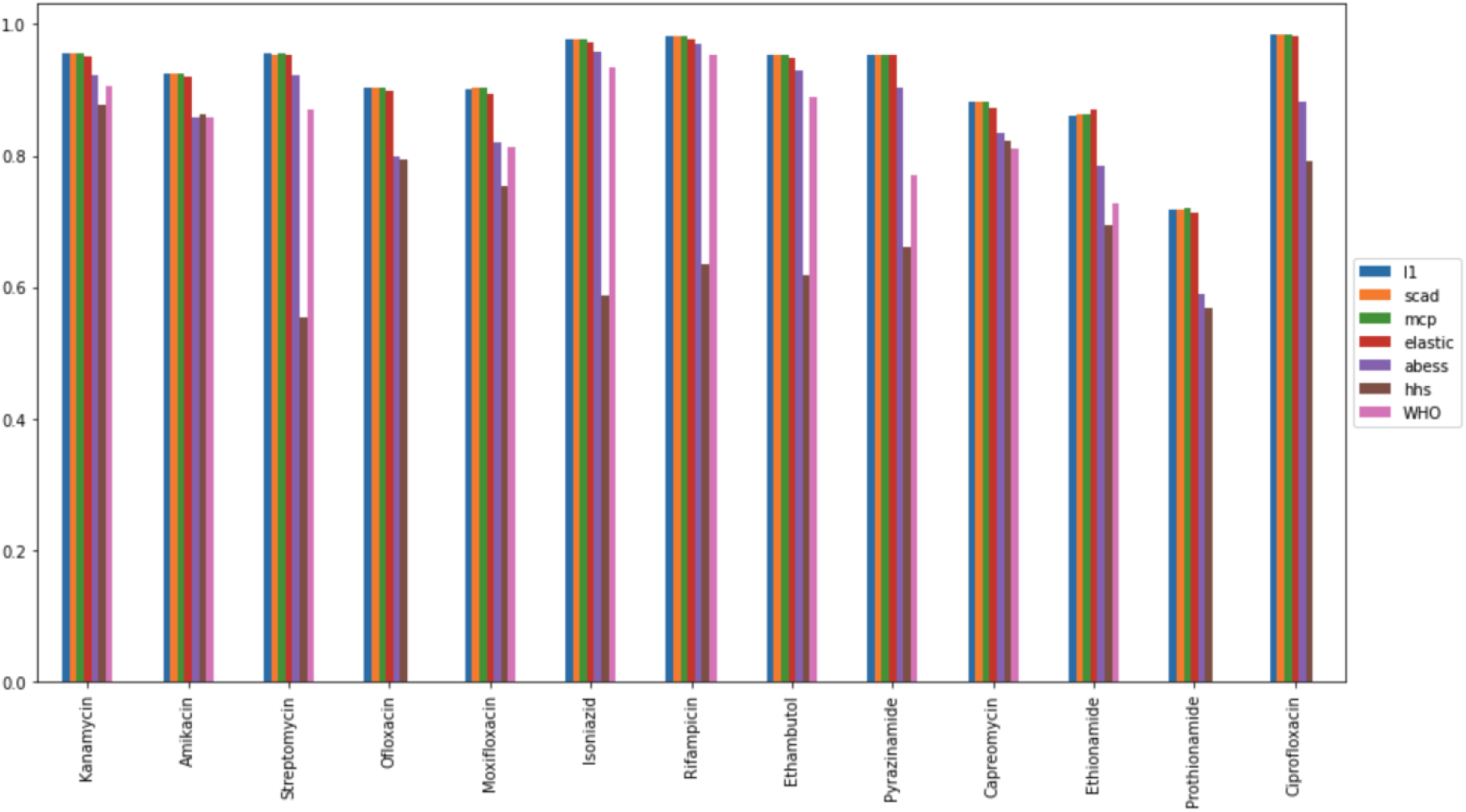
Average ROC-AUC score of logistic regression models trained on features majorly selected by regression with L1 (blue), MCP (green) and SCAD (orange) regularization, elastic net (red), HHS (brown), ABESS (purple) and WHO catalog (pink). The feature selection models were trained on training parts of 5 dataset splits and features selected on at least 3 parts were chosen. The logistic regression was trained using these features on the training partition and AUC was estimated on the testing partition. The AUC scores were averaged over all dataset splits.

We also compared the number of selected features and the stability of selection by different methods (Table S24). The numbers of selected features vary between the algorithms: logistic regression with different regularization techniques typically pick more features than ABESS and HHS. We compared the number of features selected in at least one dataset split with the number of majorly selected features (selected in at least three splits): surprisingly elastic net demonstrated the largest fraction of features selected in 3 and more splits, thus demonstrating relatively high stability, but these sets are still too large to interpret, while the second leader - ABESS - majorly selects the small number of features. (Table S24).

Next, we examined the correctness of selected features. Assuming that most of the phenotype could be explained by mutations in a relatively small number of genes that are involved in actions of drugs we compared the methods on their ability to correctly identify known associations of drugs to genes. Below, we refer to a gene as a “gene associated with a given drug” if this gene was associated with resistance to given drug in previous studies (Table 2). We refer to a gene as a “gene selected by the method for a given drug” if at least one mutation in this gene was majorly selected by this method for this drug. We compared methods by their ability to select for each drug features corresponding to genes whose associations with resistance with this particular drug are known and avoid selecting features associated with other drugs. The first comparison criterion is the Jaccard index (Table 3). In our case, it is the number of genes associated with a given drug selected by the method divided by the number of all genes associated with resistance to any drug selected for the particular drug. The ABESS method had the highest Jaccard index for 10 antibiotics. HHS was the best for Ethambutol and Kanamycin, logistic regression with elastic net regularization was best for Prothionamide. Features from logistic regression with L1, SCAD and MCP regularization showed the worst result.

**Table 3.** The comparison of abilities of the feature selection methods to catch up features for the genes associated with known mechanisms of drug resistance. Below, we refer to a gene as a “gene associated with a given drug” if this gene was associated with resistance to given drug in previous studies (Table 2). We refer to a gene as a “gene selected by the method for given drug” if at least one mutation in this gene was majorly selected by this method for this drug. In the first two rows the method name and the evaluation metric name are listed. Jaccard index is the number of genes, which are associated with a given drug selected by the method divided by a total number of genes selected by the method for this drug. True gene fraction is the number of genes associated with a given drug selected by the method divided by the number of genes associated with this drug. False gene fraction is a number of genes associated with the other drugs selected by the method divided by the number of genes associated with the other drugs. The rest of the rows present these metrics’ values for all drugs. Maximal values for all drugs and all metrics are marked in bold.

| Metric | Jaccard index |  |  |  |  |  | True gene fraction |  |  |  |  |  | False gene fraction |  |  |  |  |  |
| --- | --- | --- | --- | --- | --- | --- | --- | --- | --- | --- | --- | --- | --- | --- | --- | --- | --- | --- |
| Method | L1 | SCAD | MCP | Elastic | ABESS | HHS | L1 | SCAD | MCP | Elastic | ABESS | HHS | L1 | SCAD | MCP | Elastic | ABESS | HHS |
| Rifampicin | 0,01 | 0,01 | 0,01 | 0,01 | **0,13** | 0,02 | 0,50 | 0,50 | 0,50 | 0,50 | 0,50 | 0,50 | **0,38** | **0,38** | **0,38** | 0,31 | 0,04 | 0,08 |
| Isoniazid | 0,03 | 0,03 | 0,03 | 0,02 | **0,38** | 0,17 | **1,00** | 0,80 | 0,80 | 0,60 | 0,60 | 0,20 | **0,35** | 0,26 | 0,26 | 0,26 | 0,04 | 0,00 |
| Pyrazinamide | 0,02 | 0,03 | 0,03 | 0,01 | **0,09** | 0,01 | 0,33 | 0,33 | 0,33 | 0,33 | 0,33 | 0,33 | 0,32 | 0,32 | 0,32 | 0,40 | 0,04 | **0,44** |
| Ethambutol | 0,02 | 0,02 | 0,02 | 0,01 | 0,07 | **0,08** | **0,33** | **0,33** | **0,33** | 0,11 | 0,11 | 0,11 | **0,47** | **0,47** | **0,47** | **0,47** | 0,05 | 0,11 |
| Streptomycin | 0,06 | 0,06 | 0,06 | 0,03 | **0,33** | 0,13 | **1,00** | **1,00** | **1,00** | **1,00** | 0,67 | 0,67 | 0,24 | 0,24 | 0,24 | **0,28** | 0,08 | 0,16 |
| Kanamycin | 0,01 | 0,01 | 0,01 | 0,02 | 0,40 | **0,50** | 1,00 | 1,00 | 1,00 | 1,00 | 1,00 | 1,00 | 0,38 | **0,50** | **0,50** | 0,42 | 0,04 | 0,00 |
| Amikacin | 0,01 | 0,01 | 0,01 | 0,01 | **0,50** | 0,13 | **1,00** | **1,00** | **1,00** | 0,50 | 0,50 | 0,50 | **0,38** | **0,38** | **0,38** | 0,31 | 0,00 | 0,04 |
| Capreomycin | 0,01 | 0,01 | 0,01 | 0,01 | **0,33** | 0,29 | **1,00** | **1,00** | **1,00** | 0,50 | 0,50 | **1,00** | 0,42 | **0,50** | **0,50** | 0,27 | 0,04 | 0,00 |
| Ofloxacin | 0,00 | 0,00 | 0,00 | 0,02 | **0,25** | 0,08 | **1,00** | **1,00** | **1,00** | **1,00** | 0,50 | 0,50 | **0,50** | **0,50** | **0,50** | 0,46 | 0,00 | 0,08 |
| Moxifloxacin | 0,01 | 0,00 | 0,00 | 0,01 | **0,25** | 0,17 | **1,00** | **1,00** | **1,00** | 0,50 | 0,50 | 0,50 | 0,50 | **0,54** | **0,54** | 0,38 | 0,04 | 0,00 |
| Ciprofloxacin | 0,02 | 0,02 | 0,02 | 0,02 | **0,33** | **0,33** | 0,50 | 0,50 | 0,50 | 0,50 | 0,50 | 0,50 | **0,31** | **0,31** | **0,31** | 0,27 | 0,00 | 0,00 |
| Ethionamide | 0,02 | 0,02 | 0,11 | 0,04 | **0,40** | 0,00 | **0,67** | **0,67** | **0,67** | **0,67** | **0,67** | 0,00 | **0,36** | 0,32 | 0,24 | 0,32 | 0,04 | 0,04 |
| Prothionamide | 0,01 | 0,01 | 0,01 | **0,01** | 0,00 | 0,00 | **0,67** | **0,67** | **0,67** | **0,67** | 0,00 | 0,00 | **0,64** | 0,52 | 0,52 | 0,56 | 0,04 | 0,04 |

We also examine two additional metrics to emphasize the strength and weakness of each method: true gene fraction and false gene fraction. We define true gene fraction as the number of genes associated with a given drug selected by the algorithm for a given drug divided by the total number of genes associated with resistance to a given drug (Table 2). We define false gene fraction as the number of genes associated with the other drugs selected by the algorithm for the given drug divided by the total number of genes associated with other drugs (Table 2). The values of true gene fractions were similar for all methods, but they were slightly better for logistic regression methods. Whereas the false gene fraction values were also consistently higher for logistic regression methods (Table 3).

Genes that are selected by ABESS generally are in agreement with the literature (Table 2 and 4). Also, it’s worth noting that PFAM domain features are majorly selected by ABESS: about 53% of all majorly selected features are PFAM-domain features (Table 4).

**Table 4.**
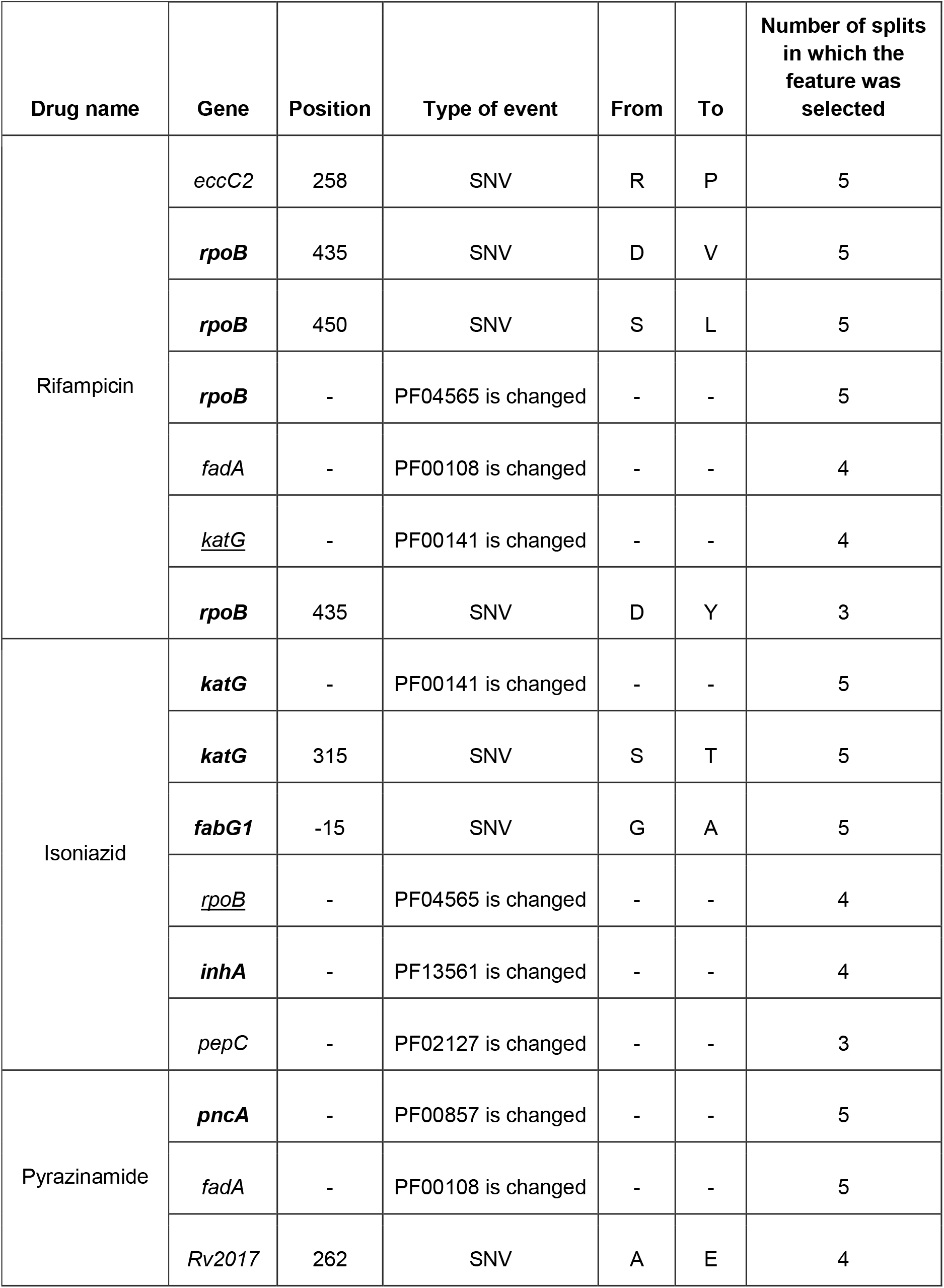
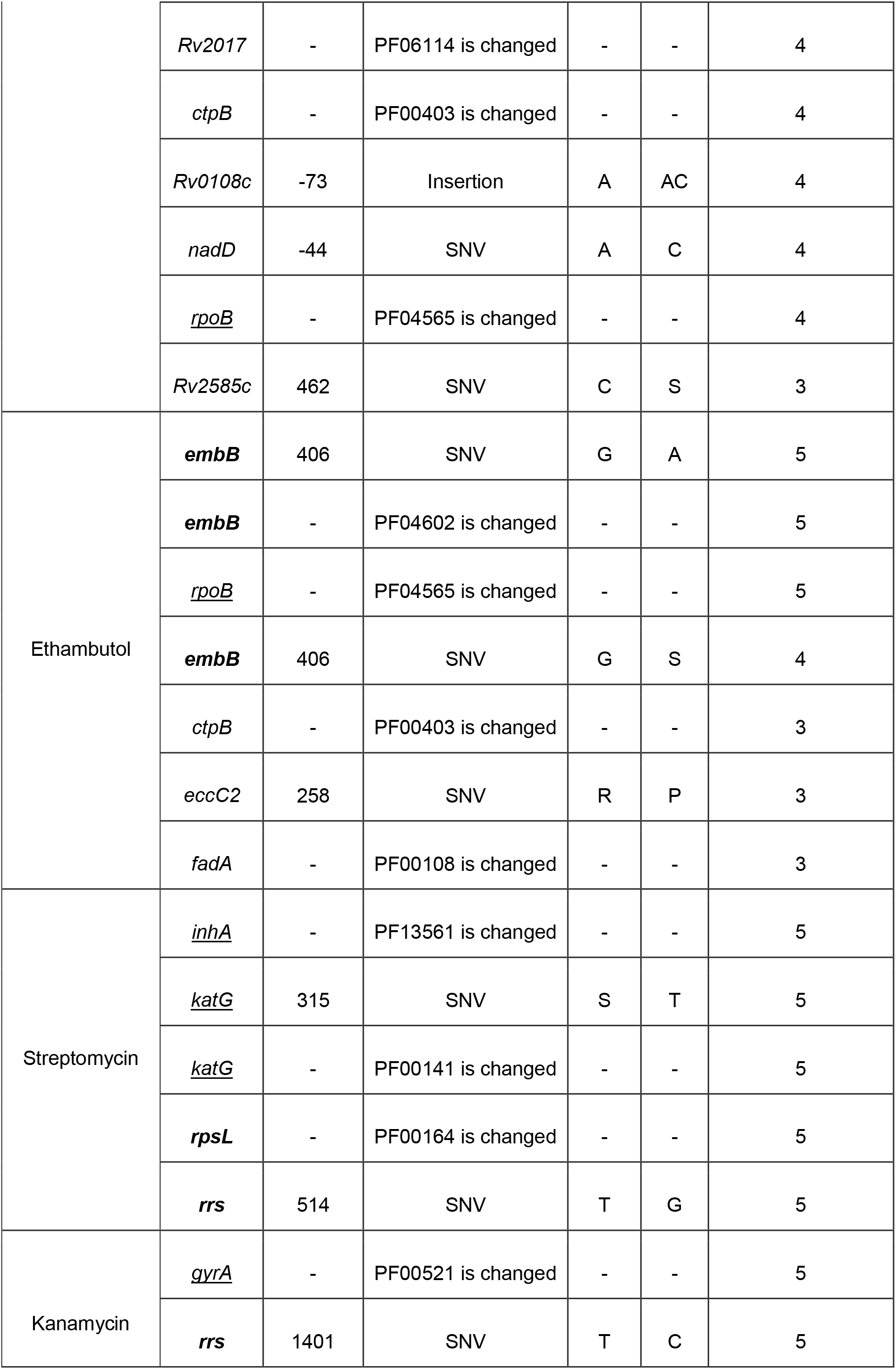
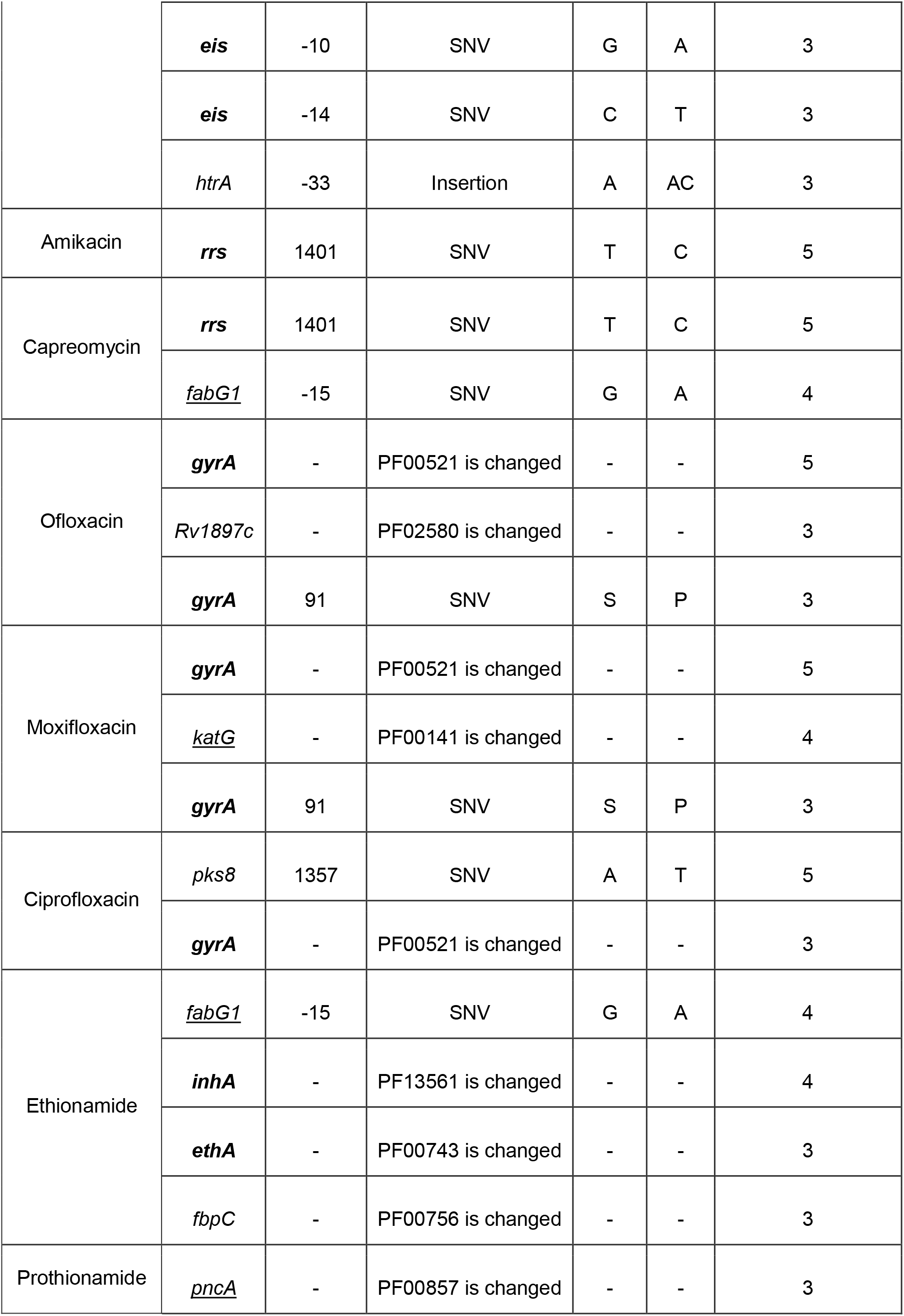
The mutations and aggregated features which are majorly selected by the ABESS algorithm. The names of genes which were associated with resistance to the given drug in previous studies (Table 2) assigned to the right drugs are marked in bold and underscored otherwise.

### ABESS is not confused by the population structure

To account for possible effects of the phylogenetic structure of the population of isolates on the results of our analysis for each drug we generated additional structural features which were corresponded to clades of the tree where prevalences of resistant isolates were different of the prevalences in the parental clades (see Methods). Each feature took values one for all isolates within the corresponded clade and zero for other isolates (see the Methods section). Structural features were generated for all drugs (see Table S25). The number of structural features varied from 1 for Prothionamide to 163 for Rifampicin (see Table S26).

To assess whether the phylogenetic features correspond to specific geographic locations or to the specific MTB phylogenetic lineages we assigned geographic locations to 3541 isolates (about 29%) for which such information was available and the information about lineages to all except two isolates, for which lineage-defining mutations from the TB Profiler table were found (Table S27). Taking together all drugs, for 139 (about 21%) of all structural features distributions of isolates by location categories were significantly different from the distributions by the same categories of all isolates having the phenotype annotations to corresponding drugs. For 480 (about 69%) of all structural features distributions of isolates by MTB phylogenetic lineages were significantly different from the distributions by categories of all isolates having the phenotype annotations to corresponding drugs (see Table S27 and Table S28). Thus, there were strong geographic and phylogenetic signals in our data.

All structural features were added to the set of genetic features for each corresponding drug and the compound feature sets were used for feature selection by the ABESS method. None of the structural features were majorly selected (Table S29), therefore, the population structure of the data did not compromise the set of features selected by ABESS.

### New associations can be found by repeating the search on the unexplained resistant isolates

Features majorly selected by ABESS did not fully explain resistance for several drugs: the significant percentage of resistant samples was not explained by the mutations selected in three or more folds, especially for the second-line drugs (Table 5). We tried to improve these results and performed the second iteration of ABESS on a reduced dataset to find additional mutations which can explain the resistance (Table 6) (see Methods). We removed from the dataset sequences with resistant phenotypes which were classified as resistant on the previous iteration. We also removed features which were majorly selected for at least one drug with positive coefficients at the first iteration enforcing ABESS to search for new associations. For six drugs the second iteration allowed to slightly increase the number of explained resistant samples (from 0.35% to 5.94%) (Table 5). Among 24 features selected at the second iteration 18 features were known from the literature as DR-associated mutations and domains in the DR-associated genes, 15 of them are correctly predicted for corresponding drugs. Two drug-associated genes which were missed at the first iteration (*embA* for Ethambutol and *gid* for Streptomycin) were found at the second iteration.

**Table 5.**
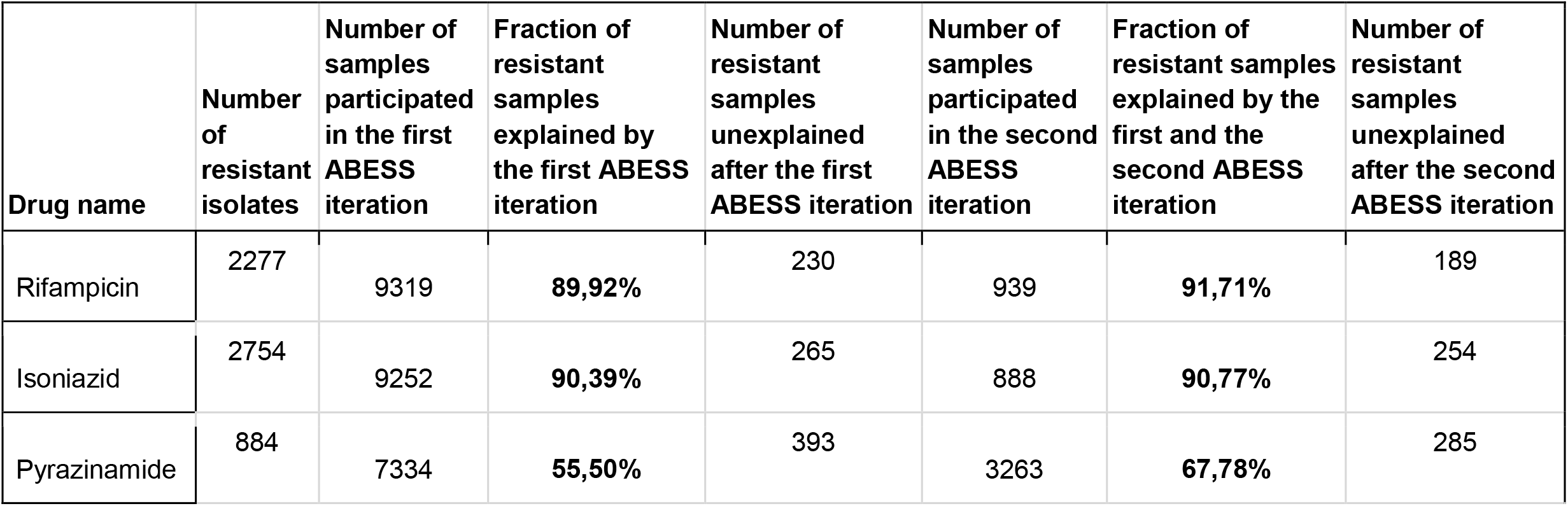
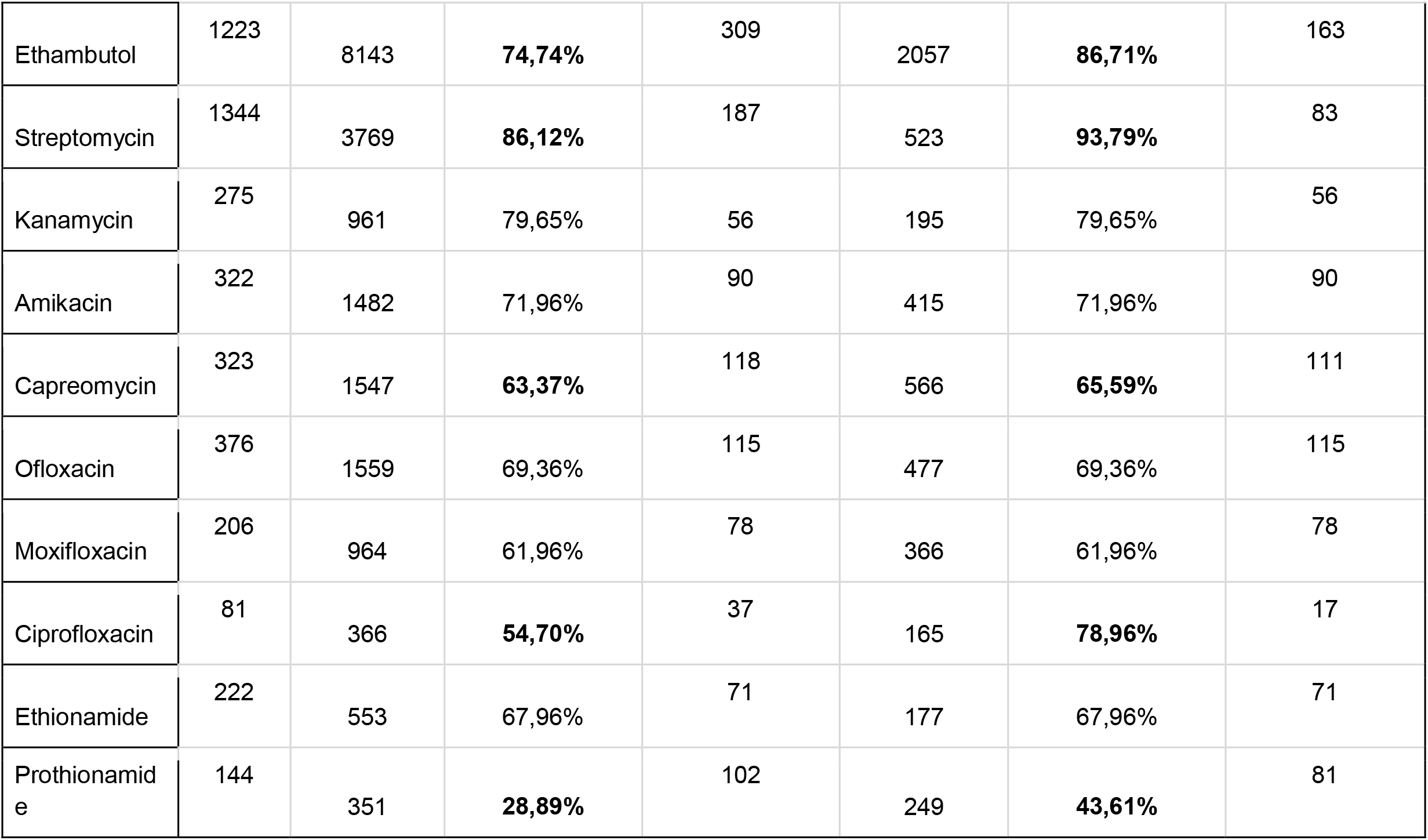
The statistics of explained resistant isolates by the first and the second iteration of the ABESS algorithm. We defined a resistant isolate to be explained by ABESS on a given dataset split if logistic regression trained on the training partition of this split using the features majorly selected by ABESS classifies this isolate as resistant. All the numbers were averaged by splits. In case of increase of the fraction of explained isolates for the given drug the values after the first and the second iteration are marked in bold.

| Drug name | Number of resistant isolates | Number of samples participated in the first ABESS iteration | Fraction of resistant samples explained by the first ABESS iteration | Number of resistant samples unexplained after the first ABESS iteration | Number of samples participated in the second ABESS iteration | Fraction of resistant samples explained by the first and the second ABESS iteration | Number of resistant samples unexplained after the second ABESS iteration |
| --- | --- | --- | --- | --- | --- | --- | --- |
| Rifampicin | 2277 | 9319 | **89,92%** | 230 | 939 | **91,71%** | 189 |
| Isoniazid | 2754 | 9252 | **90,39%** | 265 | 888 | **90,77%** | 254 |
| Pyrazinamide | 884 | 7334 | **55,50%** | 393 | 3263 | **67,78%** | 285 |
| Ethambutol | 1223 | 8143 | **74,74%** | 309 | 2057 | **86,71%** | 163 |
| Streptomycin | 1344 | 3769 | **86,12%** | 187 | 523 | **93,79%** | 83 |
| Kanamycin | 275 | 961 | 79,65% | 56 | 195 | 79,65% | 56 |
| Amikacin | 322 | 1482 | 71,96% | 90 | 415 | 71,96% | 90 |
| Capreomycin | 323 | 1547 | **63,37%** | 118 | 566 | **65,59%** | 111 |
| Ofloxacin | 376 | 1559 | 69,36% | 115 | 477 | 69,36% | 115 |
| Moxifloxacin | 206 | 964 | 61,96% | 78 | 366 | 61,96% | 78 |
| Ciprofloxacin | 81 | 366 | **54,70%** | 37 | 165 | **78,96%** | 17 |
| Ethionamide | 222 | 553 | 67,96% | 71 | 177 | 67,96% | 71 |
| Prothionamide | 144 | 351 | **28,89%** | 102 | 249 | **43,61%** | 81 |

**Table 6.**
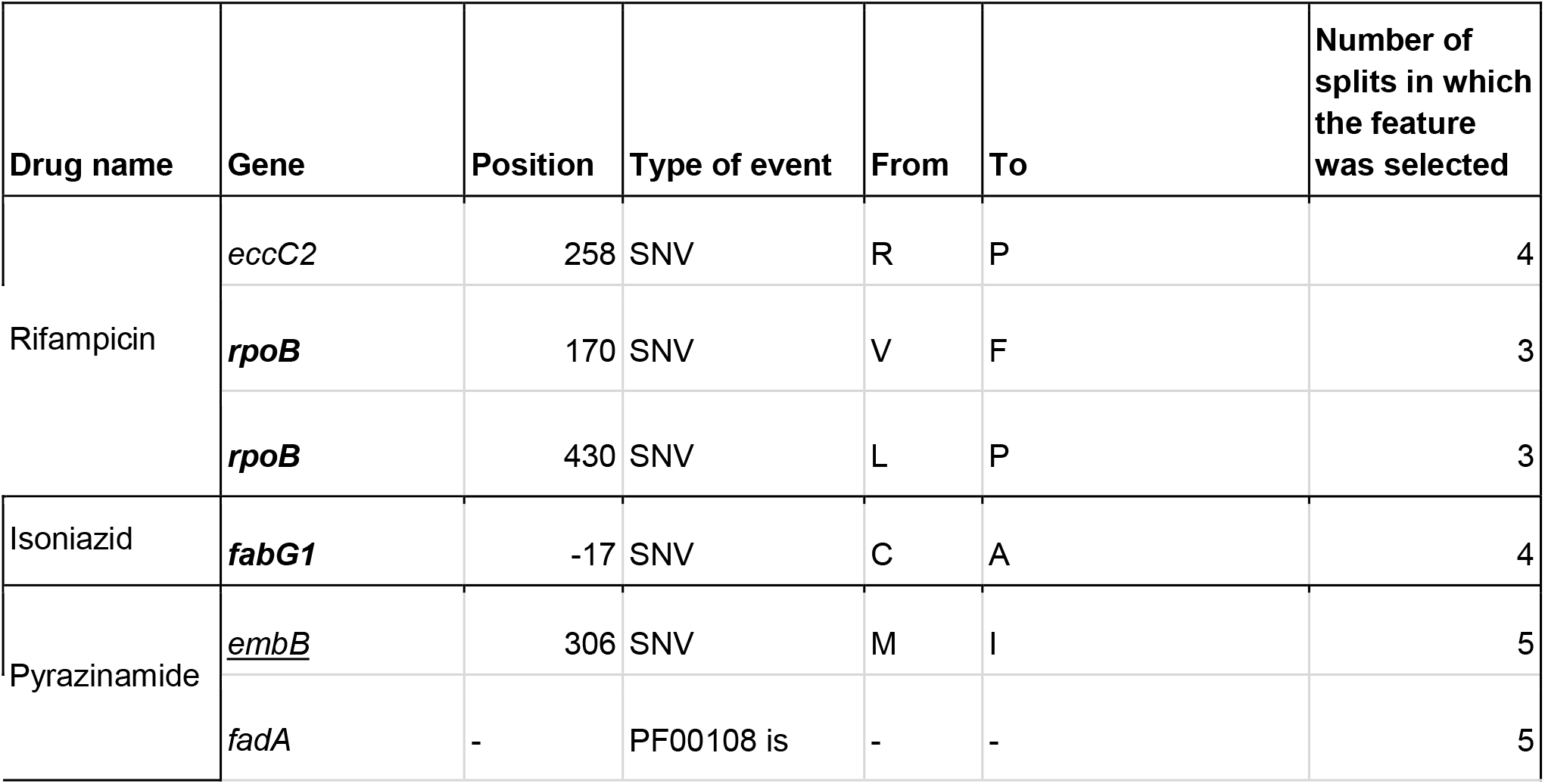
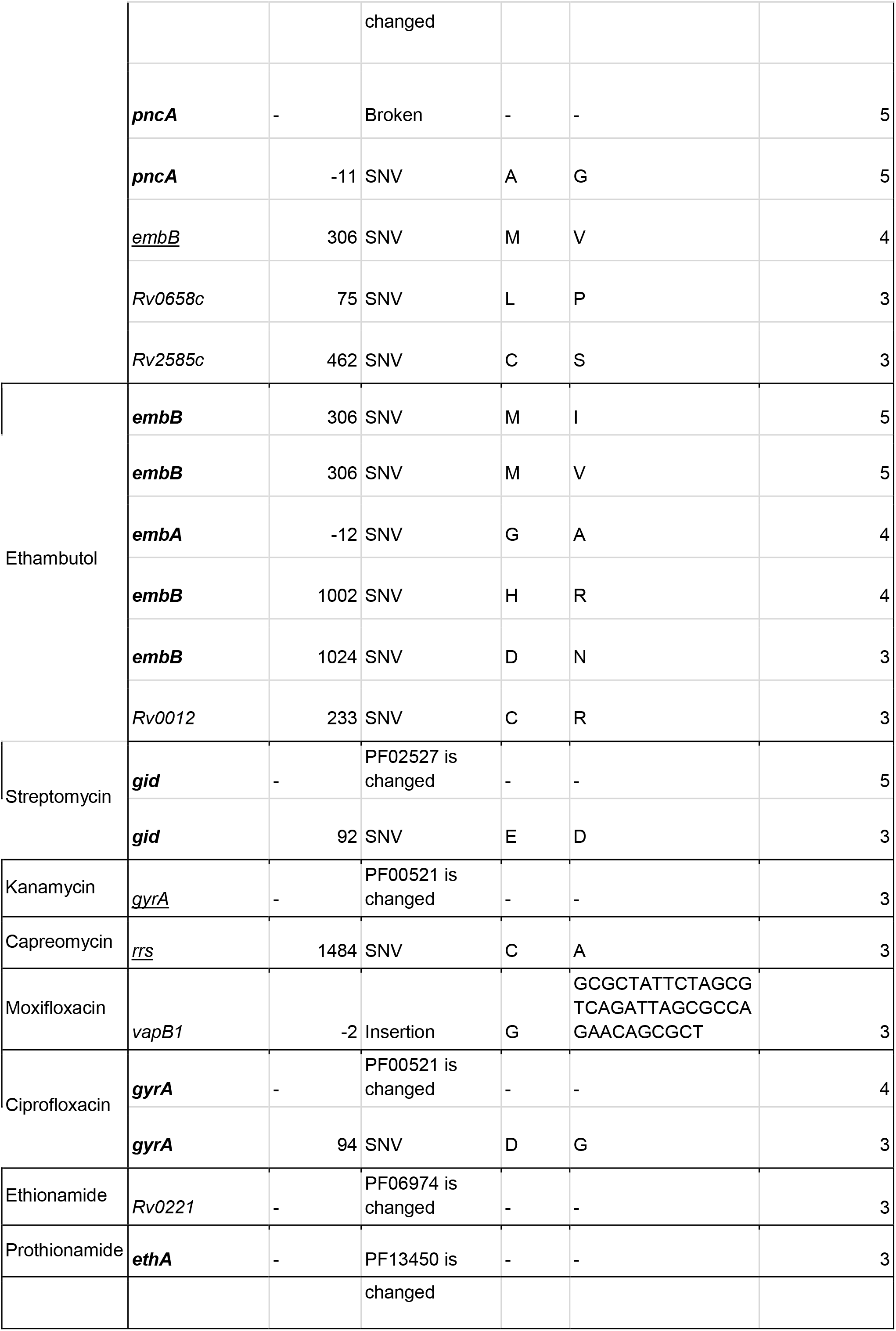
The additional mutations majorly selected by ABESS on the second iteration. The pairs of drugs and genes which were associated with each other in previous studies (Table 2) are marked in bold, while the pairs in which the gene was associated with another drug are underscored.

We tried to apply this approach to the other methods being tested above. All variants of logistic regression turned out to majorly select so many features, that these features explained all resistant samples on the first iteration. This comes at the cost of mixing correct genes with the ones associated with the other drugs according to the literature (see Jaccard Index and False gene percentage in Table 3). Meanwhile, HHS, being the worst in ROC AUC (Fig 1), explains much less resistant samples on average compared to ABESS. The percentage of explained resistant samples increased significantly after the second iteration for all drugs, but these values were lower than those for ABESS for all drugs except Pyrazinamide, Kanamycin, Amikacin, Capreomycin and Ethionamide (Table 5, Table S31). The second iteration allowed HHS to select useful mutations: from 43 features majorly selected on the second iteration 19 were point mutations and 19 were change of domain events, 5 of which were in DR-associated genes known from the literature (Table S32). Twenty of these 43 features were in the genes that had been selected on the first iteration and one of these features was a domain event; 15 domain events were selected in the new genes. Three of domain events were in known drug associated genes: the PF01751 PFAM domain change event in the *gyrB* gene was selected for Ofloxacin, the PF00743 PFAM domain change event in the *ethA* was selected for Ethionamide and the PF00141 PFAM domain change event in *katG* was selected for Isoniazid. The rpsL_K88R was selected for Streptomycin. Thus, the second iteration allowed HHS to find *gyrB, katG, rpsL and ethA*, which were not selected at the first iteration. However, compared to ABESS HHS tends to select more features and genes which were not associated with any drug in the literature.

## Discussion

We analyzed 12333 full genome MTB samples for which drug resistance status was known for at least one of 13 antibiotics being studied. For most of the samples, drug resistance status was known only for the first-line drugs. Typically, if a sample is resistant to one of the first line drugs it is resistant to the others due to stepwise acquisition of resistance-conferring mutations during long ineffective treatment by multiple drugs [64, 65]. This co-resistance decreases the power of statistical methods to find new genes and pathways weakly associated with resistance. Even the best of the algorithms being studied according to our metrics, ABESS, was unable to completely untangle cross-associations of features to different drugs (Table 4): *rpoB* is selected for Isoniazid, Ethambutol and Pyrazinamide while *katG* is selected for Rifampicin. Also, if a sample is resistant to some second-line drug it is typically resistant to one or more first-line drugs. Again, such co-resistance leads to cross associations of genes of first and second line drugs: *katG* was selected for Streptomycin and Moxifloxacin, *pncA* was selected for Prothionamide, *fabG1* was selected for Capreomycin and Ethionamide. Cross-associations between second-line drug features are also observed: the change of *gyrA* domain is selected for Kanamycin.

ABESS is the most appropriate method among the tested to solve the problem of correlation, yet at the same time, it has provided the least amount of novel genes involved in drug resistance. Totally, 16 novel genes are selected as markers of any drug resistance: 12 of them are identified after the first iteration and 4 more genes are found after the second iteration of ABESS. Some of the findings are likely to be feasible.

*eccC2* is predicted to contribute to Ethambutol and Rifampicin resistance (R258P is selected by ABESS). EccC2 belongs to the ESX-2 type VII secretion system and may be involved in virulence of the pathogen and its survival under hypoxic conditions [66].

*fadA* is associated with Ethambutol, Rifampicin and Pyrazinamide resistance (alteration of N-terminal domain is selected as an important resistance predictor). *fadA* encodes a possible acyl-CoA thiolase involved in lipid degradation and its expression is shown to be altered upon Isoniazid exposure as a compensatory response [67]. *fadA* was also associated with Ofloxacin and Kanamycin resistance in one of the early GWAS studies in MTB [1].

The metal-binding domain of *ctpB* is selected as probable Ethambutol and Pyrazinamide resistance determinant. CtpB is a plasma membrane copper (I) transporting P-type ATPase. *ctpB* transcription was activated during hypoxia, nitrosative and oxidative stress which are prevalent in the host phagosomes environment [68].

*pks8* is associated with Ciprofloxacin resistance. *pks8* encodes a probable polyketide synthase and is involved in fatty acids biosynthesis process [69]. Other polyketide synthases, *pks2* and *pks12*, are shown to be associated with MDR in other studies [70].

*fbpC* is found to be associated with Ethionamide resistance and its product is secreted antigen 85-C. Proteins of the antigen 85 complex are responsible for the high affinity of mycobacteria to fibronectin. FbpC possesses a mycolyltransferase activity required for the biogenesis of trehalose dimycolate (cord factor), a dominant structure necessary for maintaining cell wall integrity [71, 72].

Mutations in *pepC* and *htrA* are selected by ABESS as markers of resistance to Isoniazid and Kanamycin, respectively. Both of these gene products are proteases, *pepC* is shown to be expressed during long pellicle growth [73].

SNP in *nadD* upstream region is associated with Pyrazinamide resistance. NadD is involved in NAD biosynthesis, which is targeted by Isoniazid, Pyrazinamide and some other promising antimycobacterial drugs [74].

Among 4 genes selected after the second iteration of ABESS 2 genes (*Rv0012* and *Rv0658c*) probably encode conserved membrane proteins, *Rv0221* is possible triacylglycerol synthase involved in lipid metabolism and *vapB1* is an antitoxin belonging to the VapBC family. *vapB1* upstream insertion is selected as Moxifloxacin resistance marker. VapB1 is shown to be degraded in unfavorable conditions which causes toxin VapC1 release and subsequent inhibition of cell growth [75]. VapC21, a toxin from the same family, was associated with XDR in other studies [37].

Overall, novel genes selected by ABESS are likely to be compensatory and actually associated with survival in the host and drug tolerance.

HHS identified 107 novel genes after the first iteration and 21 genes after the second iteration. After additional filtering by Fisher p-value (p-value < 0.05) 44 and 1 genes remained for two iterations, correspondingly. 24 of them are associated with resistance to Pyrazinamide. There is no overlap in the selected sets of novel genes between HHS and ABESS. However, HHS output also contains some interesting findings.

HHS method recovers *rpoC* association with Streptomycin and Rifampicin resistance. Interestingly, associated variants are not the same for these drugs: single-residue replacement in site 402 of *rpoC* is selected as Rifampicin resistance marker while Streptomycin resistance is associated with replacement in site 516 of the same gene. Mutations in several sites in *rpoC* were shown to have a compensatory role for Rifampicin resistance [76] and Q1126K replacement in *rpoC* was beneficial in Rifampicin and Streptomycin double-resistant but not in sensitive strains [16].

*rpoA*, another RNA polymerase subunit, variant is associated with Pyrazinamide resistance. However, this variant (V183G) is also known to be compensatory for Rifampicin resistance [76].

Novel genes set for Pyrazinamide also contains 4 membrane transport proteins: *cycA* (involved in D-alanine, D-serine and glycine transport), *dppC* (probable ABC transporter of dipeptide across the membrane), *sugI* (involved in sugar transport) and *cysT* (probable sulfate-transport ABC transporter) [72]. *cycA* and *sugI* variants were previously associated with D-cycloserine resistance, the drug which is commonly used to treat MDR and XDR-TB [77, 78].

*nusG* variant is associated with Rifampicin resistance. NusG is a probable transcription antitermination protein which interacts with rho factor and RNA polymerase. Thus, the selected variant is likely to be compensatory for *rpoB* mutations and *nusG* regulatory mutation was previously shown to be compensatory in Rifampicin and Streptomycin double-resistant genotypes [16].

*rplU*, which encodes 50S ribosomal protein L21, is associated with Streptomycin resistance while another ribosomal protein, RpsL, and ribosomal RNA *rrs* are known to be the Streptomycin targets.

*pntAa* is also found to be associated with Streptomycin resistance and functions as a proton pump suggesting its possible role in MDR acquisition.

Logistic regression methods selected the largest set of novel genes (more than 500 novel genes remained after raw p-value filtering for each regularization technique). These large sets lack biological interpretation proving this group of methods is not suitable for the task discussed, despite the fact that these methods provide the highest prediction quality (Fig. 3).

The prediction quality of GWAS algorithms is limited by the quality of datasets. One of the reasons is samples infected by multiple strains which may lead to errors both in genotyping and phenotyping. Assigning discrete phenotypes is problematic in case of intermediate levels of drug resistance: resistant isolates with a low growth rate are interpreted as susceptible in some cases [79]. Differences in culturing and measurement protocols lead to inconsistency in phenotyping and lead to problems with the reproducibility of results. There is growing evidence that MICs are more informative than binary labels for GWAS analysis [80].

Overall, our study demonstrates the usefulness of advanced feature selection combined with feature generation by aggregation with subsequent filtering by mutation frequency. Aggregated features were majorly selected and constituted the majority of selected features. Another advantage of the aggregation is the possibility to take into account the effects of rare mutations without including them directly into the model increasing both the robustness and training speed. Also, advanced feature selection helps to treat correlations between phenotypes: simpler methods tend to explain the resistance to one drug with the mutations in the genes associated with the other drugs. Our selection strategy does not suffer from the population structure bias either: despite the strong geographic signal in our data no structure associated features are selected by the best algorithm.

## Abbreviations

MTB: *Mycobacterium tuberculosis*
DR: drug resistance
MDR: multidrug resistant
XDR: extensively drug resistant
WHO: World Health Organization
NGS: Next Generation Sequencing
PCR: polymerase chain reaction
GWAS: genome-wide association study
DA: direct association
ML: machine learning
PCA: principal component analysis
HMM: hidden Markov model
MCP: minimax concave penalty
SCAD: smoothly clipped absolute deviation
HHS: ‘hungry-hungry SNPs’
ABESS: polynomial-time algorithm for best-subset selection problem

## Authors contributions

Reshetnikov K.O.: Conceptualisation, Methodology, Software, Investigation, Writing – Original Draft Preparation, Visualization

Bykova D.I.: Conceptualisation, Methodology, Software, Investigation, Writing – Original Draft Preparation

Kuleshov K.V.: Methodology, Data Curation, Writing – Review and Editing

Chukreev K.: Software, Investigation

Guguchkin E.P.: Investigation

Akimkin V.G.: Project Administration, Funding

Neverov A.D.: Conceptualisation, Methodology, Writing – Original Draft Preparation, Writing – Review and Editing, Supervision, Project Administration

Fedonin G.G.: Conceptualisation, Methodology, Software, Investigation, Data Curation, Writing – Original Draft Preparation, Supervision

## References

1. Zhang H, Li D, Zhao L, Fleming J, Lin N, et al. Genome sequencing of 161 *Mycobacterium tuberculosis* isolates from China identifies genes and intergenic regions associated with drug resistance. Nat Genet 2013;45:1255–1260.

2. Farhat MR, Shapiro BJ, Kieser KJ, Sultana R, Jacobson KR, et al. Genomic analysis identifies targets of convergent positive selection in drug-resistant Mycobacterium tuberculosis. Nat Genet 2013;45:1183–1189.

3. Coll F, Phelan J, Hill-Cawthorne GA, Nair MB, Mallard K, et al. Genome-wide analysis of multi- and extensively drug-resistant Mycobacterium tuberculosis. Nat Genet 2018;50:307–316.

4. Yang C, Luo T, Shen X, Wu J, Gan M, et al. Transmission of multidrug-resistant Mycobacterium tuberculosis in Shanghai, China: a retrospective observational study using whole-genome sequencing and epidemiological investigation. Lancet Infect Dis 2017;17:275–284.

5. Farhat MR, Freschi L, Calderon R, Ioerger T, Snyder M, et al. GWAS for quantitative resistance phenotypes in Mycobacterium tuberculosis reveals resistance genes and regulatory regions. Nat Commun 2019;10:2128.

6. Kouchaki S, Yang Y, Walker TM, Sarah Walker A, Wilson DJ, et al. Application of machine learning techniques to tuberculosis drug resistance analysis. Bioinformatics 2019;35:2276–2282.

7. Casali N, Nikolayevskyy V, Balabanova Y, Harris SR, Ignatyeva O, et al. Evolution and transmission of drug-resistant tuberculosis in a Russian population. Nat Genet 2014;46:279–286.

8. Pankhurst LJ, del Ojo Elias C, Votintseva AA, Walker TM, Cole K, et al. Rapid, comprehensive, and affordable mycobacterial diagnosis with whole-genome sequencing: a prospective study. Lancet Respir Med 2016;4:49–58.

9. Walker TM, Kohl TA, Omar SV, Hedge J, Del Ojo Elias C, et al. Whole-genome sequencing for prediction of Mycobacterium tuberculosis drug susceptibility and resistance: a retrospective cohort study. Lancet Infect Dis. Epub ahead of print June 2015. DOI: 10.1016/S1473-3099(15)00062-6.

10. Johnsen CH, Clausen PTLC, Aarestrup FM, Lund O. Improved Resistance Prediction in Mycobacterium tuberculosis by Better Handling of Insertions and Deletions, Premature Stop Codons, and Filtering of Non-informative Sites. Front Microbiol;10. Epub ahead of print 2019. DOI: 10.3389/fmicb.2019.02464.

11. World Health Organisation. Global Tuberculosis Report; 2020.

12. Jnawali HN, Ryoo S. First– and Second–Line Drugs and Drug Resistance. In: Tuberculosis - Current Issues in Diagnosis and Management. 2013. Epub ahead of print 2013. DOI: 10.5772/54960.

13. Walker TM, Miotto P, Köser CU, Fowler PW, Knaggs J, et al. The 2021 WHO catalogue of Mycobacterium tuberculosis complex mutations associated with drug resistance: a genotypic analysis. Lancet Microbe 2022;3:e265–e273.

14. Hang NTL, Hijikata M, Maeda S, Thuong PH, Ohashi J, et al. Whole genome sequencing, analyses of drug resistance-conferring mutations, and correlation with transmission of Mycobacterium tuberculosis carrying katG-S315T in Hanoi, Vietnam. Sci Rep 2019;9:15354.

15. Trauner A, Liu Q, Via LE, Liu X, Ruan X, et al. The within-host population dynamics of Mycobacterium tuberculosis vary with treatment efficacy. Genome Biol 2017;18:71.

16. Sousa JM de, Balbontín R, Durão P, Gordo I. Multidrug-resistant bacteria compensate for the epistasis between resistances. PLOS Biol 2017;15:e2001741.

17. Andrews JM. Determination of minimum inhibitory concentrations. J Antimicrob Chemother 2001;48:5–16.

18. Tomasicchio M, Theron G, Pietersen E, Streicher E, Stanley-Josephs D, et al. The diagnostic accuracy of the MTBDRplus and MTBDRsl assays for drug-resistant TB detection when performed on sputum and culture isolates. Sci Rep 2016;6:17850.

19. Boehme CC, Nabeta P, Hillemann D, Nicol MP, Shenai S, et al. Rapid molecular detection of tuberculosis and rifampin resistance. N Engl J Med 2010;363:1005–1015.

20. Meaza A, Kebede A, Yaregal Z, Dagne Z, Moga S, et al. Evaluation of genotype MTBDRplus VER 2.0 line probe assay for the detection of MDR-TB in smear positive and negative sputum samples. BMC Infect Dis 2017;17:280.

21. Chen ML, Doddi A, Royer J, Freschi L, Schito M, et al. Beyond multidrug resistance: Leveraging rare variants with machine and statistical learning models in Mycobacterium tuberculosis resistance prediction. EBioMedicine 2019;43:356–369.

22. Pankhurst LJ, Del Ojo Elias C, Votintseva AA, Walker TM, Cole K, et al. Rapid, comprehensive, and affordable mycobacterial diagnosis with whole-genome sequencing: a prospective study. Lancet Respir Med 2016;4:49–58.

23. Walker TM, Cruz ALG, Peto TE, Smith EG, Esmail H, et al. Tuberculosis is changing. Lancet Infect Dis 2017;17:359–361.

24. The CRyPTIC Consortium and the 100,000 Genomes Project. Prediction of *Susceptibility to First-Line Tuberculosis Drugs by DNA Sequencing*. N Engl J Med 2018;379:1403–1415.

25. World Health Organization. The use of next-generation sequencing technologies for the detection of mutations associated with drug resistance in Mycobacterium tuberculosis complex: technical guide. Geneva: World Health Organization. https://apps.who.int/iris/handle/10665/274443 (2018).

26. Catalogue of mutations in Mycobacterium tuberculosis complex and their association with drug resistance. https://www.who.int/publications-detail-redirect/9789240028173 (accessed 14 August 2022).

27. Domínguez J, Boettger EC, Cirillo D, Cobelens F, Eisenach KD, et al. Clinical implications of molecular drug resistance testing for Mycobacterium tuberculosis: a TBNET/RESIST-TB consensus statement. Int J Tuberc Lung Dis 2016;20:24–42.

28. Miotto P, Tessema B, Tagliani E, Chindelevitch L, Starks AM, et al. A standardised method for interpreting the association between mutations and phenotypic drug resistance in Mycobacterium tuberculosis. Eur Respir J 2017;50:1701354.

29. Farhat MR, Sultana R, Iartchouk O, Bozeman S, Galagan J, et al. Genetic Determinants of Drug Resistance in Mycobacterium tuberculosis and Their Diagnostic Value. Am J Respir Crit Care Med 2016;194:621–630.

30. Ghosh A, N. S, Saha S. Survey of drug resistance associated gene mutations in Mycobacterium tuberculosis, ESKAPE and other bacterial species. Sci Rep 2020;10:8957.

31. Schleusener V, Köser CU, Beckert P, Niemann S, Feuerriegel S. Mycobacterium tuberculosis resistance prediction and lineage classification from genome sequencing: comparison of automated analysis tools. Sci Rep 2017;7:46327.

32. Hicks ND, Yang J, Zhang X, Zhao B, Grad YH, et al. Clinically prevalent mutations in Mycobacterium tuberculosis alter propionate metabolism and mediate multidrug tolerance. Nat Microbiol 2018;3:1032–1042.

33. Hicks ND, Carey AF, Yang J, Zhao Y, Fortune SM. Bacterial Genome-Wide Association Identifies Novel Factors That Contribute to Ethionamide and Prothionamide Susceptibility in Mycobacterium tuberculosis. mBio. Epub ahead of print 23 April 2019. DOI: 10.1128/mBio.00616-19.

34. Sergeev RS, Kavaliou IS, Sataneuski UV, Gabrielian A, Rosenthal A, et al. GenomeWide Analysis of MDR and XDR Tuberculosis from Belarus: Machine-Learning Approach. IEEE/ACM Trans Comput Biol Bioinform 2019;16:1398–1408.

35. Niehaus KE, Walker TM, Crook DW, Peto TEA, Clifton DA. Machine learning for the *prediction of antibacterial susceptibility in Mycobacterium tuberculosis*. In: IEEE-EMBS International Conference on Biomedical and Health Informatics (BHI). 2014. pp. 618–621.

36. Lees JA, Mai TT, Galardini M, Wheeler NE, Horsfield ST, et al. Improved Prediction of Bacterial Genotype-Phenotype Associations Using Interpretable Pangenome-Spanning Regressions. mBio 2020;11:e01344–20.

37. Kavvas ES, Catoiu E, Mih N, Yurkovich JT, Seif Y, et al. Machine learning and structural analysis of Mycobacterium tuberculosis pan-genome identifies genetic signatures of antibiotic resistance. Nat Commun 2018;9:1–9.

38. Deelder W, Christakoudi S, Phelan J, Benavente ED, Campino S, et al. Machine Learning Predicts Accurately Mycobacterium tuberculosis Drug Resistance From Whole Genome Sequencing Data. Front Genet;10. Epub ahead of print 2019. DOI: 10.3389/fgene.2019.00922.

39. Earle SG, Wu C-H, Charlesworth J, Stoesser N, Gordon NC, et al. Identifying lineage effects when controlling for population structure improves power in bacterial association studies. Nat Microbiol 2016;1:16041.

40. Zabeti H, Dexter N, Safari AH, Sedaghat N, Libbrecht M, et al. INGOT-DR: an interpretable classifier for predicting drug resistance in M. tuberculosis. Algorithms Mol Biol 2021;16:17.

41. Palmer AC, Chait R, Kishony R. Nonoptimal Gene Expression Creates Latent Potential for Antibiotic Resistance. Mol Biol Evol 2018;35:2669–2684.

42. Yang Y, Niehaus KE, Walker TM, Iqbal Z, Walker AS, et al. Machine learning for classifying tuberculosis drug-resistance from DNA sequencing data. Bioinformatics 2018;34:1666–1671.

43. Yang Y, Walker TM, Walker AS, Wilson DJ, Peto TEA, et al. DeepAMR for predicting co-occurrent resistance of Mycobacterium tuberculosis. Bioinformatics 2019;35:3240–3249.

44. Cohen KA, Abeel T, Manson McGuire A, Desjardins CA, Munsamy V, et al. Evolution of Extensively Drug-Resistant Tuberculosis over Four Decades: Whole Genome Sequencing and Dating Analysis of Mycobacterium tuberculosis Isolates from KwaZulu-Natal. PLoS Med 2015;12:e1001880–e1001880.

45. Manson AL, Cohen KA, Abeel T, Desjardins CA, Armstrong DT, et al. Genomic analysis of globally diverse Mycobacterium tuberculosis strains provides insights into the emergence and spread of multidrug resistance. Nat Genet 2017;49:395–402.

46. Van Rie Annelies, Warren Robin, Mshanga Idris, Jordaan Annemarie M, van der Spuy Gian D., et al. Analysis for a Limited Number of Gene Codons Can Predict Drug Resistance of Mycobacterium tuberculosis in a High-Incidence Community. J Clin Microbiol 2001;39:636–641.

47. Hazbón Manzour Hernando, Brimacombe Michael, Bobadilla del Valle Miriam, Cavatore Magali, Guerrero Marta Inírida, et al. Population Genetics Study of Isoniazid Resistance Mutations and Evolution of Multidrug-Resistant Mycobacterium tuberculosis. Antimicrob Agents Chemother 2006;50:2640–2649.

48. Burkina M, Nazarov I, Panov M, Fedonin G, Shirokikh B. Inductive Matrix Completion with Feature Selection. Comput Math Math Phys 2021;61:719–732.

49. Richardson A. Logistic Regression: A Self-Learning Text, Third Edition by David G. Kleinbaum, Mitchel Klein. Int Stat Rev 2011;79:296–296.

50. Tibshirani R. Regression Shrinkage and Selection via the Lasso. J R Stat Soc Ser B Methodol 1996;58:267–288.

51. Hoerl AE, Kennard RW. Ridge Regression: Biased Estimation for Nonorthogonal Problems. Technometrics 1970;12:55–67.

52. Hoerl AE, Kennard RW. Ridge Regression: Applications to Nonorthogonal Problems. Technometrics 1970;12:69–82.

53. Grogan TR, Elashoff DA. A simulation based method for assessing the statistical significance of logistic regression models after common variable selection procedures. Commun Stat Simul Comput 2017;46:7180–7193.

54. Saber MM, Shapiro BJ. Benchmarking bacterial genome-wide association study methods using simulated genomes and phenotypes. Microb Genomics 2020;6:e000337.

55. Fan J, Li R. Variable Selection via Nonconcave Penalized Likelihood and its Oracle Properties. J Am Stat Assoc 2001;96:1348–1360.

56. Cun-Hui Zhang. Nearly unbiased variable selection under minimax concave penalty. Ann Stat 2010;38:894–942.

57. Kumar A, Bhattacharyya S, Bouchard K. Numerical Characterization of Support Recovery in Sparse Regression with Correlated Design. http://arXiv.org/abs/ (2021).

58. Zhu J, Wen C, Zhu J, Zhang H, Wang X. A polynomial algorithm for best-subset selection problem. Proc Natl Acad Sci 2020;117:33117.

59. Libiseller-Egger J, Phelan J, Campino S, Mohareb F, Clark TG. Robust detection of point mutations involved in multidrug-resistant Mycobacterium tuberculosis in the presence of co-occurrent resistance markers. PLOS Comput Biol 2020;16:e1008518.

60. Gan GL, Nguyen MH, Willie E, Rezaie MH, Lee B, et al. Geographic heterogeneity impacts drug resistance predictions in Mycobacterium tuberculosis. 2021;2020.09.17.301226.

61. Ansari MA, Didelot X. Bayesian Inference of the Evolution of a Phenotype Distribution on a Phylogenetic Tree. Genetics 2016;204:89–98.

62. Cohen KA, Manson AL, Desjardins CA, Abeel T, Earl AM. Deciphering drug resistance in Mycobacterium tuberculosis using whole-genome sequencing: progress, promise, and challenges. Genome Med;11. Epub ahead of print December 2019. DOI: 10.1186/s13073-019-0660-8.

63. Zhang L, Zhao Y, Gao Y, Wu L, Gao R, et al. Structures of cell wall arabinosyltransferases with the anti-tuberculosis drug ethambutol. Science 2020;368:1211–1219.

64. Mnyambwa NP, Kim D-J, Ngadaya ES, Kazwala R, Petrucka P, et al. Clinical implication of novel drug resistance-conferring mutations in resistant tuberculosis. Eur J Clin Microbiol Infect Dis Off Publ Eur Soc Clin Microbiol 2017;36:2021–2028.

65. Hjort K, Jurén P, Toro JC, Hoffner S, Andersson DI, et al. Dynamics of Extensive Drug Resistance Evolution of Mycobacterium tuberculosis in a Single Patient During 9 Years of Disease and Treatment. J Infect Dis 2020;jiaa625.

66. Bose T, Das C, Dutta A, Mahamkali V, Sadhu S, et al. Understanding the role of interactions between host and Mycobacterium tuberculosis under hypoxic condition: an in silico approach. BMC Genomics 2018;19:555.

67. Abo-Kadoum M a., Dai Y, Asaad M, Hamdi I, Xie J. Differential Isoniazid Response Pattern Between Active and Dormant Mycobacterium tuberculosis. Microb Drug Resist 2021;27:768–775.

68. León-Torres A, Arango E, Castillo E, Soto CY. CtpB is a plasma membrane copper (I) transporting P-type ATPase of Mycobacterium tuberculosis. Biol Res 2020;53:6.

69. Dubey VS, Sirakova TD, Cynamon MH, Kolattukudy PE. Biochemical Function of msl5 (pks8 plus pks17) in Mycobacterium tuberculosis H37Rv: Biosynthesis of Monomethyl Branched Unsaturated Fatty Acids. J Bacteriol 2003;185:4620–4625.

70. Philalay JS, Palermo CO, Hauge KA, Rustad TR, Cangelosi GA. Genes Required for Intrinsic Multidrug Resistance in Mycobacterium avium. Antimicrob Agents Chemother 2004;48:3412–3418.

71. Belisle JT, Vissa VD, Sievert T, Takayama K, Brennan PJ, et al. Role of the major antigen of Mycobacterium tuberculosis in cell wall biogenesis. Science 1997;276:1420–1422.

72. Kapopoulou A, Lew JM, Cole ST. The MycoBrowser portal: a comprehensive and *manually annotated resource for mycobacterial genomes*. Tuberc Edinb Scotl 2011;91:8–13.

73. Kerns PW, Ackhart DF, Basaraba RJ, Leid JG, Shirtliff ME. Mycobacterium tuberculosis pellicles express unique proteins recognized by the host humoral response. Pathog Dis 2014;70:347–358.

74. Rohde KH, Sorci L. The Prospective Synergy of Antitubercular Drugs With NAD Biosynthesis Inhibitors. Front Microbiol 2021;11:3589.

75. Lu Z, Wang H, Zhang A, Tan Y. The VapBC1 toxin-antitoxin complex from Mycobacterium tuberculosis: purification, crystallization and X-ray diffraction analysis. Acta Crystallogr Sect F Struct Biol Commun 2016;72:485–489.

76. Comas I, Borrell S, Roetzer A, Rose G, Malla B, et al. Whole-genome sequencing of rifampicin-resistant Mycobacterium tuberculosis strains identifies compensatory mutations in RNA polymerase genes. Nat Genet 2012;44:106–110.

77. Chen JM, Uplekar S, Gordon SV, Cole ST. A Point Mutation in cycA Partially Contributes to the D-cycloserine Resistance Trait of Mycobacterium bovis BCG Vaccine Strains. PLOS ONE 2012;7:e43467.

78. Chen J, Zhang S, Cui P, Shi W, Zhang W, et al. Identification of novel mutations associated with cycloserine resistance in Mycobacterium tuberculosis. J Antimicrob Chemother 2017;72:3272–3276.

79. Bradley P, Gordon NC, Walker TM, Dunn L, Heys S, et al. Rapid antibiotic-resistance predictions from genome sequence data for Staphylococcus aureus and Mycobacterium tuberculosis. Nat Commun 2015;6:10063.

80. Ahmad S, Mokaddas E, Al-Mutairi N, Eldeen HS, Mohammadi S. Discordance across Phenotypic and Molecular Methods for Drug Susceptibility Testing of Drug-Resistant Mycobacterium tuberculosis Isolates in a Low TB Incidence Country. PLOS ONE 2016;11:e0153563.

