## Supplementary Materials for "Feature selection and aggregation for antibiotic resistance GWAS in *Mycobacterium tuberculosis*: a comparative study"

#### Overview of the feature selection methods

We have chosen to use logistic regression with L1, minimax concave penalty (MCP) and smoothly clipped absolute deviation (SCAD) regularization, elastic net, ‘Hungry, Hungry SNPos‘ (HHS), and a polynomial-time algorithm for best-subset selection problem (ABESS) in this study. Below is a brief description of these methods.

The logistic regression model searches for additive coefficients $\bar{\beta}$ of genomic variations to maximize the likelihood of the dataset - the probability to observe the dataset with given $\bar{\beta}$. Wald and likelihood statistics allow estimating statistical significance of coefficients $\bar{\beta}$ [1].

Likelihood maximization tends to assign nonzero weights to all features, underestimating the impact of causative features and overfitting the noise. Also, due to correlations with benign features, the causative ones may be lost in the process of the selection. Lasso regularization [2] adds a linear regularization term penalizing large coefficients which are also known as the L1 norm regularization. As a result, only one feature is chosen from a subset of highly correlated ones. Ridge regression [3, 4] penalizes the squares of $\bar{\beta}$ coefficients (also known as the L2 norm), while elastic net combines both penalties. Lasso was shown to outperform unregularized stepwise feature selection both in the quality of prediction and in the percentage of causative features in the selected set on simulated data [2, 5]. The lasso achieves better or even performance in simulated bacterial GWAS compared to LMM models even in the presence of high LD (strong population structure) [6]. Elastic net was earlier shown to outperform lasso in prediction quality [7].

Penalizing large coefficients comes at a price: large true coefficients will be punished. It's obvious that any regularization will have its own drawbacks and limitations, but new regularization terms are constantly being invented. Fan and Li developed SCAD regularization which has a nonlinear decreasing penalty for large coefficients [8]. A similar effect is achieved by MCP regularization [9]. MCP and SCAD feature selection outperformed lasso on all types of simulated data except for non-sparse data with a low signal-to-noise ratio, in which case the elastic net was the leader [10].

Some methods penalize the number of nonzero coefficients (L0 norm regularization). The functional being optimized is no longer differentiable: enumeration of feature subsets is required. ABESS goes further and can optimize various information criteria (AIC, BIC, cross-validation, and an original asymptotic criterion - SIC) searching for the best subset [11]. For each feature set size from 1 to the user-defined maximum, the best subset is constructed and the final subset is selected based on the given information criterion. The best subset of a given size is constructed by starting from the features which are the most correlated with the target variable and iteratively improving the solution by scoring all the features with the score dependent on the currently selected set and exchanging the worst features to the best ones. This procedure is shown to converge in a finite number of steps [11]. The authors showed that ABESS outperformed SCAD, MCP, and Lasso regularization on simulated data.

Authors also provide a deep theoretical analysis of the ABESS algorithm studying the sufficient conditions of correctness and proving that if such conditions hold the algorithm's running time is polynomial. Like LASSO, SCAD, and MCP the ABESS algorithm finds the correct model if the correlations between features are not too big as well as the true number of causative features, the true coefficients are not too small while the data is not too sparse. All the methods have difficulties when there are a lot of small effects [11].

Along with well-known and theoretically grounded approaches described above, new heuristic methods are being invented for GWAS analysis which try to cope with correlations between features by taking into account the population structure [12]. HHS algorithm starts with assigning a score reflecting the correlation with the phenotype corrected to the population structure to each SNP. The score is adjustable to the class imbalance in the phenotype and allele frequencies. Similar to ABESS the currently selected set of features (all the features with positive scores) is optimized iteratively by decreasing the normalized scores of mutually correlated features in the selected set until the convergence [12].

### Supplementary Methods

#### Dataset and raw data processing

The dataset contains 13258 whole-genome sequences of *M. tuberculosis* for which both phenotypes and raw data (Illumina short reads) were available from the literature [13–22]. TruSeq3-PE-2 adapters and low-quality ends (Q<30 with sliding window of length 10) were trimmed using trimmomatic v.0.38 (java -Xmx1800m -jar trimmomatic-0.38.jar PE -phred33 R1.fastq.gz R2.fastq.gz out_p1.fastq.gz out_u1.fastq.gz out_p2.fastq.gz out_u2.fastq.gz ILLUMINACLIP:TruSeq3-PE-2.fa:1:30:10 SLIDINGWINDOW:10:30). Next, raw reads were mapped onto the reference genome H37Rv (version 3) by bwa mem with default parameters and resulting bam files were sorted and indexed with samtools. Then we marked duplicates using MarkDuplicates for the Picard package. In cases when multiple runs corresponded to one biosample we merged such bam files in one after the previous step.

Variant calling was performed with HaplotypeCaller from the GATK suite (version 4) [23]. Variants were called jointly using GenomicsDBImport and GenotypeGVCFs tools: at first, gvcf files were produced with HaplotypeCaller, then multiple gvcfs were imported to GenomicsDB and, finally, the db was jointly genotyped with GenotypeGVCFs producing joint vcf. Since joint calling of the whole dataset was computationally infeasible, we've split the dataset to three subsets, genotyped all the subsets jointly and then split joint vcfs to individual ones. After that, multiple filtering stages were performed.

At first, the variants were filtered by quality with recommended parameters (hard filtering with QD >= 2, FS <=60, MQ >= 40, MQRankSum >= -12.5, ReadPosRankSum >= -8, SOR <= 3). After that only variants covered with at least 10 reads (DP >= 10) and having allele frequency higher than 75% were keeped. Variants from regions annotated as “Repeat_region” or “mobile_element” according to Mycobrowser v.3 annotation were also removed.

At the second stage both variants and samples were filtered with iterative procedure: we first delete all variants, which are covered with less than 10 reads in less than 90% of the samples, from all the samples; then we delete all samples in which less than 90% of variable loci (preserved on previous iteration) are covered with at least 10 reads. This was repeated until convergence. This left us with 12354 sequences for further analysis.

Next step was the annotation based on Mycobrowser v.3. To take into account possible start codon loss or appearance of the inframe stop codons along with frameshifts and correctly annotate SNVs, insertions and deletions, we've performed a special translation procedure. For each isolate we took the nucleotide sequence of each protein-coding gene along with its 180 bp upstream and downstream regions which differed from the reference sequence and ran blastx [24] to compare this nucleotide sequence with the reference protein sequence of a given gene. Next, the first start codon upstream of the first position of the best blastx hit among all reading frames was found. The isolate's nucleotide sequence was translated from this start codon to the first stop codon within the same reading frame. The protein sequence obtained was aligned to the reference protein sequence with Needleman–Wunsch algorithm implemented in biopython [25]. The gene was labeled as “broken” if it was shorter than half the reference protein length or 30% longer than the reference protein. If the gene is broken, no mutation is considered in this gene. Otherwise, all mutations inside the gene are annotated according to position in protein based on the alignment. SNVs, insertions and deletions at a distance less than 100 bp upstream of a gene were annotated as upstream relative to this gene. Nucleotide variants in noncoding genes were annotated using the reference genome gene boundaries. Synonymous mutations were excluded from the further analysis.

#### Splitting dataset for further experiments

The final dataset was split to 13 smaller datasets containing only samples with known phenotypes for each antibiotic. Each sample is typically a part of multiple datasets. All feature generation, parameter tuning, feature selection and quality estimation procedures were based on 5-fold cross-validation for each drug separately. For each drug the dataset was randomly split into four non overlapping subsets (folds). Each fold was used as the testing set one at a time, while the union of the others served as the training set. Below we call every such partitioning “the dataset split”. For every dataset split the testing fold is called “the testing partition” and the union of the others is called “the training partition”.

#### Aggregation of mutations

We examined three types of aggregated features: an indicator of the presence of any mutation in a gene, an indicator that the gene is broken, i.e. it has a loss of function mutation, and features indicating alternation of PFAM domains.

If there is any mutation in a gene then the indicator of the presence of mutations in this gene takes the value of “1” and “0” otherwise. We call this approach “gene aggregation”. “Broken gene” is another binary feature. Gene is considered to be ‘broken’ if the length of its open reading frame (ORF) is decreased due to mutations by half or increased by 30% relative to the annotated ORF length (see Dataset and raw data processing). The design of “the domain X has been changed” features is described in more detail in the next section.

After the aggregated feature generation the mutations and the other binary features that had happened less than 3 times were excluded from the dataset. Then, the features whose values fully coincide, i.e. having correlation equal to 1, on the training partition of the given dataset split were unified and were treated as one feature during the selection process. For presentation purposes the same numerical ID was assigned to all features constituting unified ones.

#### PFAM domains based feature generation

Gene names from the Mycobrowser v.3 were used to get domains’ markup and pretrained HMM models from PFAM database (PFAM-A 33.1 version). For each sample, HMM scores were computed for the mutated sequences of the domains by using the maximum a posteriori method. The python module “pomegranate” (v. 0.12.2) was used to implement a scoring function for trained HMM models: the score was a logistic probability that the trained model generated a given sequence.

To move from continuous HMM scores to binary features for each drug and each domain and each dataset split we found a threshold for HMM score. The isolates from the training partition were ordered by the Δscore - an absolute difference between an HMM score of the domain of an isolate and a corresponding HMM score of the H37Rv domain sequence. The threshold was selected to split the dataset maximizing the difference of frequencies of susceptible and resistant isolates:

$$threshold = argmax_{threshold}(f_S(\Delta score \geq threshold) - f_R(\Delta score \geq threshold))$$

where, $f_S$(Δscore ≥ threshold) is a frequency of susceptible isolates with a Δscore more or equal to the threshold, $f_R$(Δscore ≥ threshold) is an analogous frequency of resistant isolates.

Accordingly, for each isolate and each drug and each dataset split we labeled a domain with HMM score deviating from the wild-type score more or equal to the threshold found as “changed” so that an additional type of binary features “the domain X has been changed” appeared.

The source code for generating of PFAM domain features is available on github: https://github.com/Reshetnikoff/m.tuberculosis-research-code

### Training of the feature selection algorithms and their performance benchmarking

The python implementation of regularized logistic regression (LR) from the Picasso package [26] was used in our study for SCAD, MCP and L1 regularization and scikit-learn (v. 0.22.1) was used for LR with elastic net regularization. The feature selection procedure by these algorithms was simply the gathering of features having nonzero coefficients on the training partition.

The ABESS implementation v. 0.1.0 from the original paper [11] with default parameters (family=’binomial’, normalize=0, max_splicing_iter=30, warm.start=False, tune.type=’gic’, num.threads=24) was used.

For every dataset split the training partition was used for the training of the selection algorithms. After the training, a list of features that have non-zero coefficients in the trained model was obtained. Logistic regression (LR) (with solver=’saga’, max_iter=10000, without regularization) from the “scikit-learn” package (v. 0.22.1) was trained using these features on the same training partition . The testing partition was used for assessing the prediction quality of the selected features using the ROC-AUC metric and F1 score.

To estimate the statistical significance of the differences in performance of the models trained on the feature sets comprising various aggregated features (see the “Estimating the impact of aggregation” in the main text) the procedure described above was repeated 100 times and Wilcoxon signed-rank test was used from the scipy python module to compute pairwise p-values (Table S2-S7,S9-10).

#### Searching for hyperparameter values

SCAD, MCP and elastic net have two hyperparameters which tune regularization. The first hyperparameter of LR with elastic net regularization from the scikit-learn package is C, the inverse of regularization strength, the second is l1_ratio, which regulates the relative importance of L1 and L2 regularization. SCAD and MCP both have lambda and alpha parameters: to understand their meaning see the papers [27, 28].

We selected three or four arbitrary significantly different values of the second parameter for parametrization of LR regularization functions : l1_ratio equaled 0.25, 0.5 and 0.75 for elastic net and alpha equaled 3, 5, 10 and 15 for SCAD and MCP. For each selected value of the second parameter we found the minimal value of the first parameter (1/C and lambda respectively) for which all features have zero regression coefficients. After that, the interval from the zero value to the found value was divided into 20 equal parts. The value in the middle of each interval and the corresponding value of the second parameter formed pairs of hyperparameter values which were used for estimating the ROC AUC as follows.

For each pair of hyperparameter values the corresponding LR models were trained on the training partition. The algorithm contains two steps: the feature selection step and the assessment step of selected features. On the feature selection step the features having nonzero coefficients were selected. On the next assessment step, logistic regression (solver='saga', max_iter=10000, without regularization) was trained on the training partition using features selected on the feature selection step and then used for predictions of dependent variables (drug phenotypes) for samples from the testing partition. The pairs of hyperparameters corresponding to the highest roc-auc metrics averaged over five dataset splits were choused. This procedure was applied separately for each drug. Finally, for each drug the best pair of hyperparameter values chosen for each LR regularization function was used for further analysis.

Logistic regression with L1 regularization has only one hyperparameter. It was tuned in the same way as lambda for SCAD and MCP, as it described above.

The original implementation of HHS v. 0.2.0 was used in this study [12]. The method has one hyperparameter - the minimum frequency of mutations that lead to the resistance phenotype. The values of the hyperparameter equal to 1, 3 and 5 were tested. For each hyperparameter value and for each drug HHS algorithm was used for the feature selection. Then, selected features were used for training of the LR model (solver='saga', max_iter=10000, without regularization) on the training partition of each dataset split. For each drug, this parameter was selected to achieve the best prediction performance measured as average roc-auc on test partitions of dataset splits.

#### Selection of the final majorly selected feature set

After the training, a list of features that have non-zero coefficients in each trained model was obtained for each dataset split. For each feature from the lists the number of splits in which this feature was selected was computed (Table S11-S16). We defined a feature to be majorly selected if it was selected in three or more dataset splits (Table 4). For each feature (including domain features), Fisher exact test p-value was computed using the “scipy” python package on the testing partition of the first dataset split. This allowed us to evade the overfitting bias of domain aggregation feature binarization: the threshold was obtained on the training partition, while the p-value was calculated on the whole dataset. This procedure allowed us to filter by p-value the majorly selected features for the subsequent search for their biological interpretation. Unfortunately, some of the stably selected features were rare enough to never occur in the test partition of that particular split, so these features were excluded from this analysis. Finally, for ABESS and HHS the majorly selected features were filtered by Fisher p-value with threshold 0.05. We also report the mean values of coefficients of all selected features from the trained models (Table S11-S16).

### Accounting for the population structure

The phylogenetic tree was built with IQ-Tree (v. 2.2.0) using the GTR+G model. Since full genomes are too large the pseudo alignment was used instead of full MSA: all constant positions and indels were excluded. The positions in the genes associated with the drug resistance in the literature (Table 2) were also excluded.

The big tree was pruned for each drug using the “prune” function from the ete3 (v. 3.0) python package [29] keeping only leaves with known phenotype. TreeBreaker v. 1.1 was used to find the clades on each phylogenetic tree with phenotype distribution different from the parental clades. The resulting annotated trees were parsed with ete3 and internal nodes with posterior probability of distribution switch higher than 0.5 were selected.

The locations of the samples were obtained from the NCBI BioSample database using the Entrez module from the biopython (v. 1.79) package [25].

The correspondence of samples to the TB-Profiler phylogenetic lineages were inferred using the lineage-defining SNPs from the TB-Profiler [30] database (https://github.com/jodyphelan/TBProfiler/tree/master/db/tbdb.barcode.bed version from 2021.01.11). Then for each sample, the lineages were sorted from higher order lineages to their sublineages and only the deepest sublineage was kept.

### The second iteration of ABESS and HHS

We defined a resistant isolate to be explained by ABESS on a given dataset split if logistic regression trained on the training partition of this split using the features majorly selected by ABESS classifies this isolate as resistant. The fraction of explained isolates was computed for all isolates in the training partition of each dataset split and values obtained for all splits were averaged. The coefficients of the logistic regression models trained on all splits were averaged for each selected feature.

The second iteration was performed on the same dataset splits. For each split all resistant isolates explained on the first iteration were removed. Then random susceptible isolates were removed to preserve the fraction of the resistant isolates in the dataset. All features which were majorly selected for at least one drug with positive coefficients at the first iteration were completely removed from the dataset at the second iteration enforcing ABESS to search for new associations. After the ABESS training feature selection process was repeated: we computed the number of times each feature was selected. Then logistic regression was trained again on the set of majorly selected features, the regression coefficients were averaged and new explained isolates were counted for each drug (Table 5,6 and S30). The same procedure was performed for HHS algorithm (Table S31-S33)

### Supplementary figures

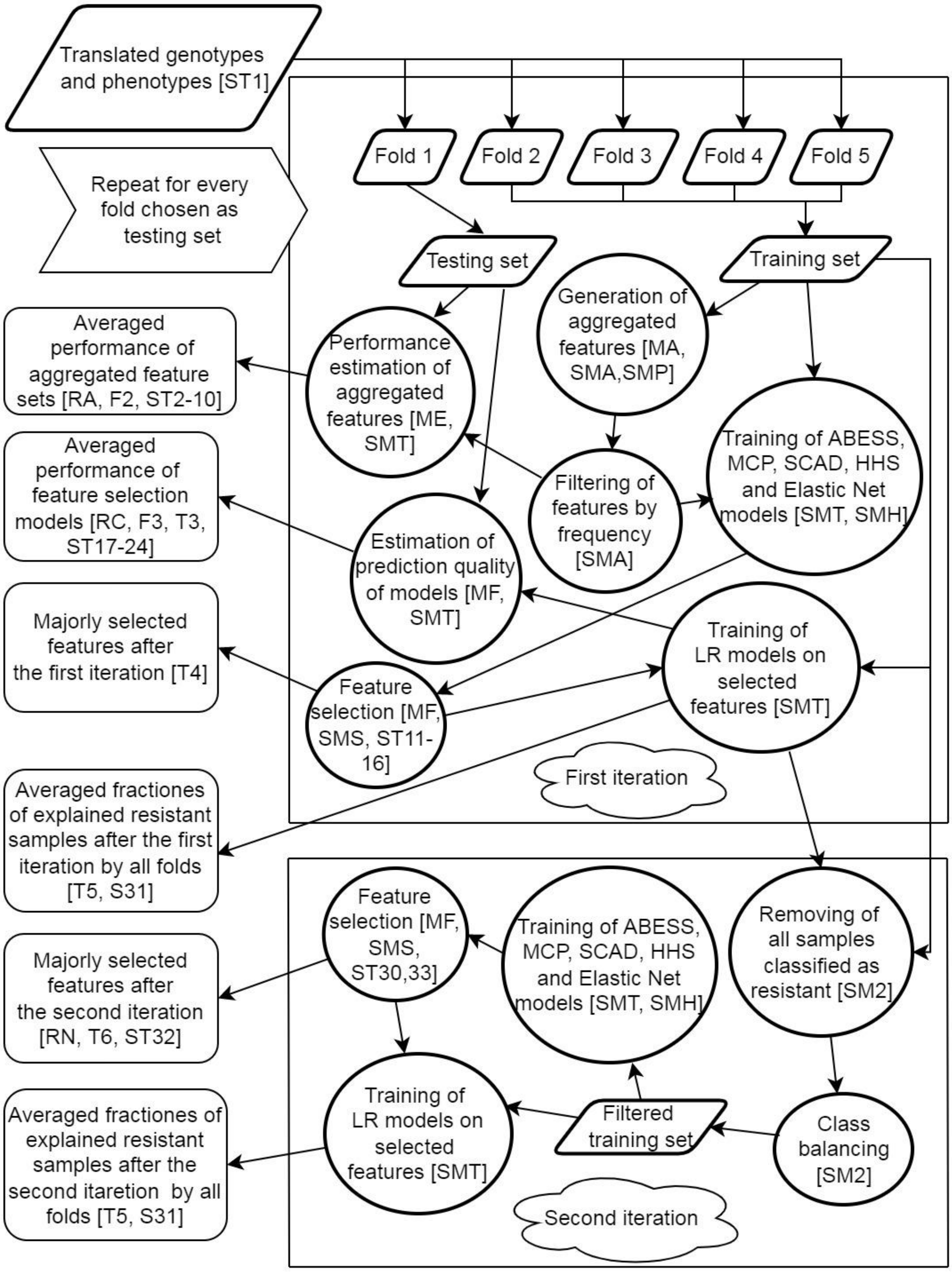


Fig. S1. The main steps of the study. To read about each step see the section listed in the square brackets (i.e. [MA] etc.). In Methods: MA - “Aggregation of mutations”, ME - “Estimating the impact of aggregation”, MF - “Feature selection and model quality evaluation”; in Supplementary Methods: SMA - “Aggregation of mutations”, SMP - “PFAM domains based feature generation”, SMT - “Training of the feature selection algorithms and their performance benchmarking”, SMH - “Searching for hyperparameter values”, SMS - “Selection of the final majorly selected feature set”, SM2 - “The second iteration of ABESS and HHS”; in Results: RA - “Aggregation of mutations improves the prediction quality”, RC - “Comparison of the feature selection algorithms”, RN - “New associations can be found by repeating the search on the unexplained resistant isolates”. F# means Figure #, T# means Table #, ST# means Supplementary Table #.

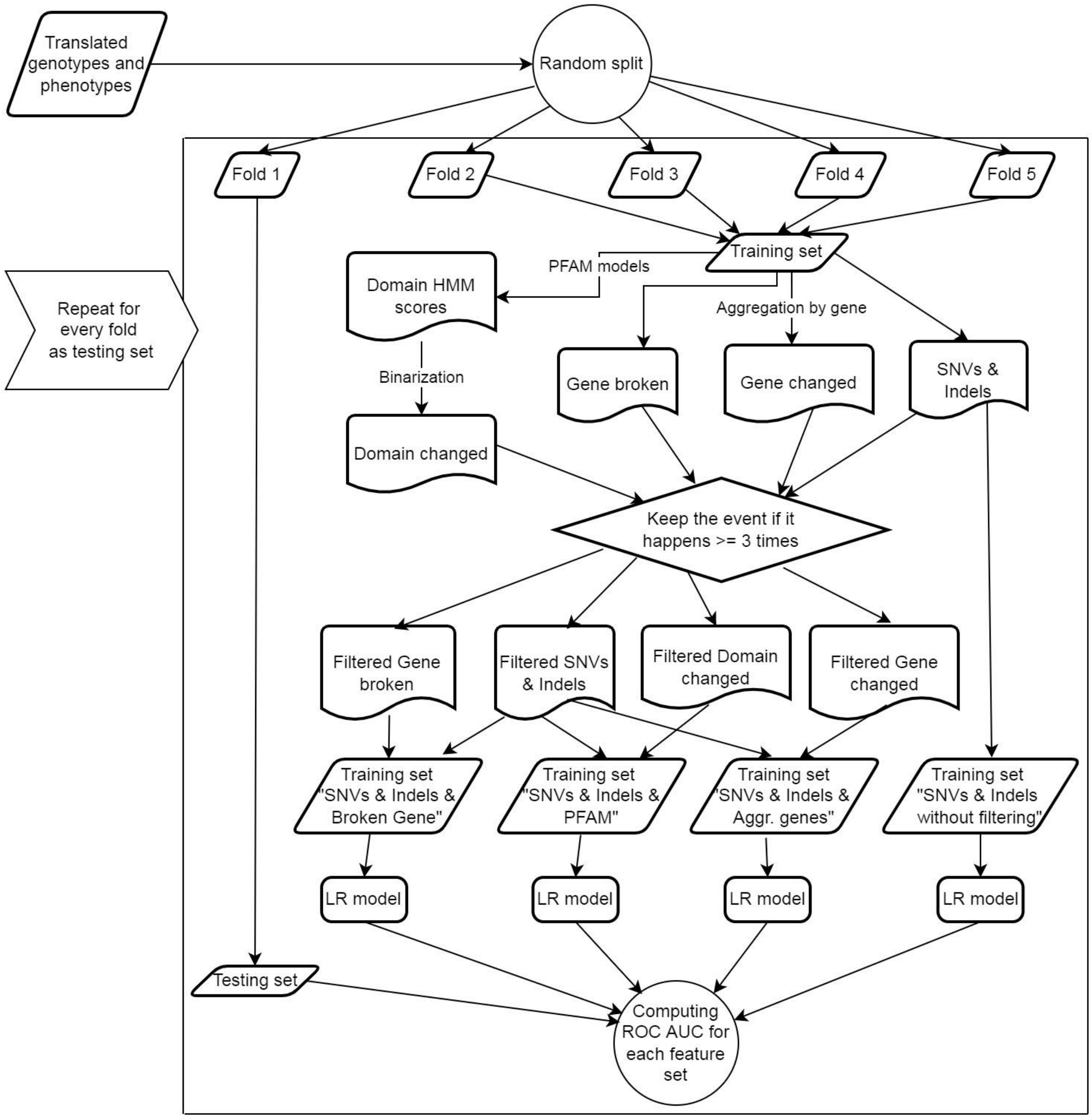


Fig. S2. The detailed schema of the comparison of feature aggregation strategies.

### Supplementary tables.

Table S1. Experimentally validated phenotypes of all isolates from our dataset with data sources. "S" means susceptible, "R" means resistant, "-" means unknown. The columns represent drugs for which the phenotypes are known for at least one isolate from the dataset. The last column contains the referent to the source publication.

https://github.com/Reshetnikoff/m.tuberculosis-research-code/blob/main/Supplementary%20tables/Table%20S1.zip

Table S2. The mean ROC AUC of logistic regression estimated by multiple 5 fold cross-validations for “SNPs and Indels” and “SNPs, Indels and PFAM domain features excluding rare mutations” feature sets. The p-value is calculated by the Wilcoxon signed-rank test on 100 random replications. Higher values for each drug and group are marked in bold.

| | **SNPs & Indels** | **SNPs & Indels & PFAM** | **P-value** |
|---|---|---|---|
| Rifampicin | **0,9825** | 0,9815 | 7,1E-18 |
| Isoniazid | **0,9759** | 0,9756 | 1,2E-06 |
| Pyrazinamide | 0,9439 | **0,9507** | 3,9E-18 |
| Ethambutol | **0,9579** | 0,9523 | 3,9E-18 |
| Streptomycin | 0,9421 | **0,9511** | 3,9E-18 |
| Kanamycin | **0,9515** | 0,9494 | 1,2E-10 |
| Amikacin | 0,9198 | **0,9223** | 3,2E-14 |
| Capreomycin | 0,8769 | **0,8845** | 4,5E-18 |
| Ofloxacin | **0,9087** | 0,9055 | 1,3E-15 |
| Moxifloxacin | 0,8961 | 0,8956 | 4,5E-01 |
| Ciprofloxacin | 0,9866 | 0,9856 | 1,4E-02 |
| Ethionamide | 0,8236 | **0,8566** | 3,9E-18 |
| Prothionamide | 0,6976 | **0,7227** | 5,0E-18 |
| First line | 0,9605 | **0,9622** | 3,9E-18 |
| Aminoglycosides | 0,9226 | **0,9268** | 3,9E-18 |
| Fluoroquinolones | **0,9304** | 0,9289 | 1,3E-12 |
| Second line | 0,8826 | **0,8903** | 3,9E-18 |
| All drugs | 0,9126 | **0,9179** | 3,9E-18 |

Table S3. The mean ROC AUC of logistic regression estimated by multiple 5 fold cross-validations for “SNPs and Indels” and “SNPs, Indels and “broken gene” features excluding rare mutations” feature sets. The p-value is calculated by the Wilcoxon signed-rank test on 100 random replications. Higher values for each drug and group are marked in bold.

| | **SNPs & Indels** | **SNPs & Indels & Broken genes** | **P-value** |
|---|---|---|---|
| Rifampicin | **0,9825** | 0,9822 | 4,3E-14 |

| | | | |
|---|---|---|---|
| Isoniazid | **0,9759** | 0,9757 | 1,1E-11 |
| Pyrazinamide | 0,9439 | **0,9450** | 3,6E-16 |
| Ethambutol | 0,9579 | **0,9584** | 3,4E-15 |
| Streptomycin | 0,9421 | **0,9427** | 1,5E-12 |
| Kanamycin | 0,9515 | **0,9524** | 5,8E-09 |
| Amikacin | 0,9198 | **0,9210** | 5,1E-11 |
| Capreomycin | **0,8769** | 0,8738 | 1,5E-17 |
| Ofloxacin | 0,9087 | **0,9091** | 2,7E-03 |
| Moxifloxacin | **0,8961** | 0,8946 | 1,6E-08 |
| Ciprofloxacin | 0,9866 | **0,9868** | 8,2E-03 |
| Ethionamide | 0,8236 | **0,8364** | 3,9E-18 |
| Prothionamide | 0,6976 | **0,6997** | 3,9E-04 |
| First line | 0,9605 | **0,9608** | 6,9E-17 |
| Aminoglycosides | 0,9226 | 0,9225 | 2,1E-01 |
| Fluoroquinolones | **0,9304** | 0,9302 | 3,5E-03 |
| Second line | 0,8826 | **0,8843** | 3,5E-17 |
| All drugs | 0,9126 | **0,9137** | 1,4E-17 |

Table S4. The mean ROC AUC of logistic regression estimated by multiple 5 fold cross-validations for “SNPs and Indels” and “SNPs, Indels and “gene aggregation”” feature sets. The p-value is calculated by the Wilcoxon signed-rank test on 100 random replications. Higher values for each drug and group are marked in bold.

| | **SNPs & Indels** | **SNPs & Indels & gene aggregation** | **P-value** |
|---|---|---|---|
| Rifampicin | **0,9825** | 0,9810 | 3,9E-18 |
| Isoniazid | **0,9759** | 0,9748 | 1,7E-17 |
| Pyrazinamide | 0,9439 | **0,9444** | 5,4E-04 |
| Ethambutol | **0,9579** | 0,9524 | 3,9E-18 |
| Streptomycin | **0,9421** | 0,9407 | 1,2E-14 |
| Kanamycin | **0,9515** | 0,9460 | 4,5E-18 |
| Amikacin | **0,9198** | 0,9176 | 2,2E-14 |
| Capreomycin | **0,8769** | 0,8756 | 1,6E-05 |
| Ofloxacin | 0,9087 | **0,9096** | 7,9E-05 |
| Moxifloxacin | 0,8961 | **0,9024** | 5,6E-18 |
| Ciprofloxacin | **0,9866** | 0,9861 | 6,0E-02 |
| Ethionamide | 0,8236 | **0,8431** | 3,9E-18 |
| Prothionamide | **0,6976** | 0,6891 | 1,7E-14 |

| | | | |
|---|---|---|---|
| First line | **0,9605** | 0,9587 | 3,9E-18 |
| Aminoglycosides | **0,9226** | 0,9200 | 3,9E-18 |
| Fluoroquinolones | 0,9304 | **0,9327** | 2,3E-17 |
| Second line | 0,8826 | **0,8837** | 6,2E-11 |
| All drugs | 0,9126 | 0,9125 | 5,3E-01 |

Table S5. The mean F1 score of logistic regression estimated by multiple 5 fold cross-validations for “SNPs and Indels” and “SNPs, Indels and PFAM domain features excluding rare mutations” feature sets. The p-value is calculated by the Wilcoxon signed-rank test on 100 random replications. Higher values for each drug and group are marked in bold.

| | **SNPs & Indels** | **SNPs & Indels & PFAM** | **P-value** |
|---|---|---|---|
| Rifampicin | 0,9348 | **0,9379** | 5,9E-18 |
| Isoniazid | 0,9322 | **0,9362** | 3,9E-18 |
| Pyrazinamide | 0,7153 | **0,7607** | 3,9E-18 |
| Ethambutol | **0,7580** | 0,7508 | 4,1E-18 |
| Streptomycin | 0,8577 | **0,8711** | 3,9E-18 |
| Kanamycin | **0,8736** | 0,8679 | 9,9E-12 |
| Amikacin | 0,8298 | **0,8342** | 1,1E-09 |
| Capreomycin | **0,7450** | 0,7405 | 6,5E-07 |
| Ofloxacin | **0,7850** | 0,7757 | 5,8E-16 |
| Moxifloxacin | **0,7088** | 0,7017 | 4,0E-07 |
| Ciprofloxacin | 0,8926 | **0,9046** | 8,6E-13 |
| Ethionamide | 0,6684 | **0,7040** | 3,9E-18 |
| Prothionamide | 0,5555 | **0,5937** | 4,5E-18 |
| First line | 0,8396 | **0,8513** | 3,9E-18 |
| Aminoglycosides | 0,8265 | **0,8285** | 1,3E-07 |
| Fluoroquinolones | 0,7955 | 0,7940 | 5,3E-02 |
| Second line | **0,7573** | 0,7653 | 6,5E-18 |
| All drugs | **0,7890** | 0,7984 | 3,9E-18 |

Table S6. The mean F1 score of logistic regression estimated by multiple 5 fold cross-validations for “SNPs and Indels” and “SNPs, Indels and “broken gene” features excluding rare mutations” feature sets. The p-value is calculated by the Wilcoxon signed-rank test on 100 random replications. Higher values for each drug and group are marked in bold.

| | SNPs & Indels | SNPs & Indels & Broken gene | P-value |
|---|---|---|---|
| Rifampicin | **0,9348** | 0,9342 | 5,3E-09 |
| Isoniazid | 0,9322 | 0,9324 | 1,2E-02 |
| Pyrazinamide | 0,7153 | **0,7192** | 6,5E-10 |
| Ethambutol | **0,7580** | 0,7547 | 5,9E-13 |
| Streptomycin | 0,8577 | **0,8585** | 2,8E-03 |
| Kanamycin | 0,8736 | **0,8766** | 5,7E-07 |
| Amikacin | 0,8298 | 0,8306 | 5,3E-02 |
| Capreomycin | 0,7450 | 0,7439 | 3,5E-02 |
| Ofloxacin | **0,7850** | 0,7821 | 1,3E-08 |
| Moxifloxacin | **0,7088** | 0,7047 | 9,3E-05 |
| Ciprofloxacin | 0,8926 | 0,8917 | 1,2E-01 |
| Ethionamide | **0,6684** | 0,6925 | 4,8E-18 |
| Prothionamide | 0,5555 | 0,5529 | 1,2E-01 |
| First line | 0,8396 | 0,8398 | 1,7E-01 |
| Aminoglycosides | 0,8265 | **0,8274** | 1,8E-03 |
| Fluoroquinolones | **0,7955** | 0,7928 | 1,5E-08 |
| Second line | 0,7573 | **0,7594** | 4,6E-09 |
| All drugs | 0,7890 | **0,7903** | 8,3E-09 |

Table S7. The mean F1 score of logistic regression estimated by multiple 5 fold cross-validations for “SNPs and Indels” and “SNPs, Indels and “gene aggregation”” feature sets. The p-value is calculated by the Wilcoxon signed-rank test on 100 random replications. Higher values for each drug and group are marked in bold.

| | SNPs & Indels | SNPs & Indels & gene aggregation | P-value |
|---|---|---|---|
| Rifampicin | **0,9348** | 0,9340 | 7,2E-06 |
| Isoniazid | 0,9322 | 0,9320 | 2,6E-01 |
| Pyrazinamide | 0,7153 | **0,7357** | 3,9E-18 |
| Ethambutol | **0,7580** | 0,7499 | 4,3E-18 |
| Streptomycin | 0,8577 | **0,8596** | 2,4E-07 |
| Kanamycin | **0,8736** | 0,8714 | 8,8E-04 |
| Amikacin | 0,8298 | **0,8317** | 1,6E-04 |
| Capreomycin | **0,7450** | 0,7414 | 5,7E-06 |
| Ofloxacin | **0,7850** | 0,7765 | 3,3E-17 |
| Moxifloxacin | 0,7088 | **0,7131** | 6,8E-05 |
| Ciprofloxacin | 0,8926 | **0,8966** | 8,6E-04 |

| | | | |
|---|---|---|---|
| Ethionamide | 0,6684 | **0,6910** | 6,9E-18 |
| Prothionamide | 0,5555 | 0,5553 | 8,6E-01 |
| First line | 0,8396 | **0,8422** | 7,7E-16 |
| Aminoglycosides | 0,8265 | 0,8260 | 1,3E-01 |
| Fluoroquinolones | 0,7955 | 0,7954 | 6,8E-01 |
| Second line | 0,7573 | **0,7596** | 8,0E-08 |
| All drugs | 0,7890 | **0,7914** | 1,5E-13 |

Table S8. The unbiased sample variance of ROC AUC of logistic regression with L1 regularization trained on different sets of features: “SNPs and Indels”; “SNPs, Indels and PFAM domain features excluding rare mutations”; “SNPs, Iindels and “broken gene” features excluding rare mutations”; “SNPs, Indels and “gene aggregation” features excluding rare mutations”.

| | **SNPs & Indels** | **SNPs & Indels & PFAM** | **SNPs & Indels & broken genes** | **SNPs & Indels & gene aggregation** |
|---|---|---|---|---|
| Rifampicin | 3,9E-07 | 6,1E-07 | 4,6E-07 | 6,0E-07 |
| Isoniazid | 4,8E-07 | 6,8E-07 | 5,2E-07 | 6,1E-07 |
| Pyrazinamide | 2,3E-06 | 2,3E-06 | 2,7E-06 | 2,7E-06 |
| Ethambutol | 6,5E-07 | 1,2E-06 | 5,9E-07 | 9,3E-07 |
| Streptomycin | 2,7E-06 | 2,2E-06 | 3,0E-06 | 3,3E-06 |
| Kanamycin | 1,3E-05 | 8,9E-06 | 1,2E-05 | 1,3E-05 |
| Amikacin | 1,1E-05 | 1,2E-05 | 1,2E-05 | 1,3E-05 |
| Capreomycin | 2,2E-05 | 2,6E-05 | 2,0E-05 | 2,3E-05 |
| Ofloxacin | 1,1E-05 | 1,3E-05 | 1,2E-05 | 9,7E-06 |
| Moxifloxacin | 2,0E-05 | 2,9E-05 | 2,1E-05 | 2,3E-05 |
| Ciprofloxacin | 1,2E-05 | 2,4E-05 | 1,2E-05 | 1,3E-05 |
| Ethionamide | 4,4E-05 | 5,9E-05 | 5,4E-05 | 5,3E-05 |
| Prothionamide | 1,5E-04 | 1,5E-04 | 1,6E-04 | 1,8E-04 |

Table S9. The mean ROC AUC of logistic regression estimated by multiple 5 fold cross-validations for “SNPs, Indels and PFAM domain features excluding rare mutations’ and ‘SNPs, Indels and "gene aggregation" features excluding rare mutations” feature sets. The p-value is calculated by the Wilcoxon signed-rank test on 100 random replications. Higher values for each drug and group are marked in bold.

| | **SNPs & Indels & PFAM** | **SNPs & Indels & gene aggregation** | **P-value** |
|---|---|---|---|
| Rifampicin | **0,9815** | 0,9810 | 3,4E-08 |

| | | | |
|---|---|---|---|
| Isoniazid | **0,9756** | 0,9748 | 1,2E-13 |
| Pyrazinamide | **0,9507** | 0,9444 | 3,9E-18 |
| Ethambutol | 0,9523 | 0,9524 | 2,3E-01 |
| Streptomycin | **0,9511** | 0,9407 | 3,9E-18 |
| Kanamycin | **0,9494** | 0,9460 | 1,2E-15 |
| Amikacin | **0,9223** | 0,9176 | 4,1E-18 |
| Capreomycin | **0,8845** | 0,8756 | 3,9E-18 |
| Ofloxacin | 0,9055 | **0,9096** | 2,3E-17 |
| Moxifloxacin | 0,8956 | **0,9024** | 3,8E-17 |
| Ciprofloxacin | 0,9856 | 0,9861 | 3,0E-01 |
| Ethionamide | **0,8566** | 0,8431 | 5,8E-18 |
| Prothionamide | **0,7227** | 0,6891 | 3,9E-18 |
| First line | **0,9622** | 0,9587 | 3,9E-18 |
| Aminoglycosides | **0,9268** | 0,9200 | 3,9E-18 |
| Fluoroquinolones | 0,9289 | **0,9327** | 4,7E-18 |
| Second line | **0,8903** | 0,8837 | 3,9E-18 |
| All drugs | **0,9179** | 0,9125 | 3,9E-18 |

Table S10. The mean F1 score of logistic regression estimated by multiple 5 fold cross-validations for “SNPs, Indels and PFAM domain features excluding rare mutations’ and ‘SNPs, Indels and "gene aggregation" features excluding rare mutations” feature sets. The p-value is calculated by the Wilcoxon signed-rank test on 100 random replications. Higher values for each drug and group are marked in bold.

| | **SNPs & Indels & PFAM** | **SNPs & Indels & gene aggregation** | **P-value** |
|---|---|---|---|
| Rifampicin | **0,9379** | 0,9340 | 4,7E-18 |
| Isoniazid | **0,9362** | 0,9320 | 3,9E-18 |
| Pyrazinamide | **0,7607** | 0,7357 | 3,9E-18 |
| Ethambutol | 0,7508 | 0,7499 | 2,6E-02 |
| Streptomycin | **0,8711** | 0,8596 | 3,9E-18 |
| Kanamycin | 0,8679 | **0,8714** | 2,0E-05 |
| Amikacin | **0,8342** | 0,8317 | 8,3E-05 |
| Capreomycin | 0,7405 | 0,7414 | 2,6E-01 |
| Ofloxacin | 0,7757 | 0,7765 | 4,2E-01 |
| Moxifloxacin | 0,7017 | **0,7131** | 4,5E-11 |
| Ciprofloxacin | **0,9046** | 0,8966 | 5,4E-08 |
| Ethionamide | **0,7040** | 0,6910 | 4,4E-12 |

| | | | |
|---|---|---|---|
| Prothionamide | **0,5937** | 0,5553 | 1,4E-17 |
| First line | **0,8513** | 0,8422 | 3,9E-18 |
| Aminoglycosides | **0,8285** | 0,8260 | 2,6E-09 |
| Fluoroquinolones | 0,7940 | 0,7954 | 6,4E-02 |
| Second line | **0,7653** | 0,7596 | 5,9E-16 |
| All drugs | **0,7984** | 0,7914 | 4,1E-18 |

Table S11. The features selected by ABESS. The features whose values fully coincide on the given dataset split were unified and were treated as one feature during the selection process. All features were enumerated for each drug and the same IDs were assigned to the features constituting unified ones.

https://github.com/Reshetnikoff/m.tuberculosis-research-code/blob/main/Supplementary%20tables/Table%20S11.tsv

Table S12. The features selected by HHS. The features whose values fully coincide on the given dataset split were unified and were treated as one feature during the selection process. All features were enumerated for each drug and the same IDs were assigned to the features constituting unified ones.

https://github.com/Reshetnikoff/m.tuberculosis-research-code/blob/main/Supplementary%20tables/Table%20S12.tsv

Table S13. The features selected by logistic regression with SCAD regularization. The features whose values fully coincide on the given dataset split were unified and were treated as one feature during the selection process. All features were enumerated for each drug and the same IDs were assigned to the features constituting unified ones.

https://github.com/Reshetnikoff/m.tuberculosis-research-code/blob/main/Supplementary%20tables/Table%20S13.tsv

Table S14. The features selected by logistic regression with MCP regularization. The features whose values fully coincide on the given dataset split were unified and were treated as one feature during the selection process. All features were enumerated for each drug and the same IDs were assigned to the features constituting unified ones.

https://github.com/Reshetnikoff/m.tuberculosis-research-code/blob/main/Supplementary%20tables/Table%20S14.tsv

Table S15. The features selected by logistic regression with L1 regularization. The features whose values fully coincide on the given dataset split were unified and were treated as one feature during the selection process. All features were enumerated for each drug and the same IDs were assigned to the features constituting unified ones.

https://github.com/Reshetnikoff/m.tuberculosis-research-code/blob/main/Supplementary%20tables/Table%20S15.tsv

Table S16. The features selected by logistic regression with elastic net regularization. The features whose values fully coincide on the given dataset split were unified and were treated as one feature during the selection process. All features were enumerated for each drug and the same IDs were assigned to the features constituting unified ones.

https://github.com/Reshetnikoff/m.tuberculosis-research-code/blob/main/Supplementary%20tables/Table%20S16.tsv

Table S17. The quality metrics of ABESS obtained by 5-fold cross-validation.

| Drug name | AUC | Sensitivity | Specivity | NPV | PPV |
|---|---|---|---|---|---|
| Kanamycin | 0,92 | 0,77 | 0,98 | 0,91 | 0,94 |
| Amikacin | 0,86 | 0,72 | 0,99 | 0,93 | 0,96 |
| Streptomycin | 0,92 | 0,75 | 0,96 | 0,88 | 0,92 |
| Ofloxacin | 0,80 | 0,68 | 0,93 | 0,90 | 0,75 |
| Moxifloxacin | 0,82 | 0,56 | 0,94 | 0,89 | 0,72 |
| Isoniazid | 0,96 | 0,90 | 0,98 | 0,96 | 0,96 |
| Rifampicin | 0,97 | 0,90 | 0,99 | 0,97 | 0,96 |
| Ethambutol | 0,93 | 0,75 | 0,94 | 0,95 | 0,69 |
| Pyrazinamide | 0,90 | 0,62 | 0,97 | 0,95 | 0,71 |
| Capreomycin | 0,84 | 0,64 | 0,96 | 0,91 | 0,82 |
| Ethionamide | 0,78 | 0,60 | 0,89 | 0,77 | 0,78 |
| Prothionamide | 0,59 | 0,33 | 0,85 | 0,65 | 0,61 |
| Ciprofloxacin | 0,88 | 0,57 | 0,97 | 0,89 | 0,85 |

Table S18. The quality metrics of HHS obtained by 5-fold cross-validation.

| Drug name | AUC | Sensitivity | Specivity | NPV | PPV |
|---|---|---|---|---|---|
| Amikacin | 0,86 | 0,72 | 0,99 | 0,93 | 0,96 |
| Ofloxacin | 0,79 | 0,58 | 0,98 | 0,88 | 0,91 |
| Pyrazinamide | 0,66 | 0,20 | 0,99 | 0,90 | 0,83 |
| Rifampicin | 0,64 | 0,24 | 1,00 | 0,80 | 0,96 |
| Capreomycin | 0,82 | 0,66 | 0,96 | 0,91 | 0,81 |
| Ciprofloxacin | 0,79 | 0,53 | 0,98 | 0,88 | 0,89 |
| Ethambutol | 0,62 | 0,24 | 0,99 | 0,88 | 0,73 |
| Ethionamide | 0,69 | 0,54 | 0,85 | 0,74 | 0,69 |
| Isoniazid | 0,59 | 0,18 | 0,99 | 0,74 | 0,93 |
| Kanamycin | 0,88 | 0,76 | 0,98 | 0,91 | 0,95 |

| | | | | | |
|---|---|---|---|---|---|
| Moxifloxacin | 0,75 | 0,48 | 0,97 | 0,87 | 0,81 |
| Prothionamide | 0,57 | 0,18 | 0,95 | 0,63 | 0,76 |
| Streptomycin | 0,56 | 0,14 | 0,97 | 0,67 | 0,73 |

Table S19. The quality metrics of logistic regression with SCAD regularization obtained by 5-fold cross-validation.

| Drug name | AUC | Sensitivity | Specivity | NPV | PPV |
|---|---|---|---|---|---|
| Kanamycin | 0,95 | 0,83 | 0,96 | 0,93 | 0,89 |
| Amikacin | 0,92 | 0,75 | 0,98 | 0,93 | 0,92 |
| Streptomycin | 0,95 | 0,84 | 0,94 | 0,91 | 0,88 |
| Ofloxacin | 0,90 | 0,70 | 0,95 | 0,91 | 0,81 |
| Moxifloxacin | 0,90 | 0,62 | 0,95 | 0,90 | 0,77 |
| Isoniazid | 0,98 | 0,90 | 0,99 | 0,96 | 0,97 |
| Rifampicin | 0,98 | 0,91 | 0,99 | 0,97 | 0,97 |
| Ethambutol | 0,95 | 0,75 | 0,96 | 0,96 | 0,75 |
| Pyrazinamide | 0,95 | 0,74 | 0,97 | 0,96 | 0,80 |
| Capreomycin | 0,88 | 0,65 | 0,96 | 0,91 | 0,80 |
| Ethionamide | 0,86 | 0,69 | 0,86 | 0,81 | 0,77 |
| Prothionamide | 0,72 | 0,58 | 0,74 | 0,72 | 0,61 |
| Ciprofloxacin | 0,98 | 0,90 | 0,99 | 0,97 | 0,96 |

Table S20. The quality metrics of logistic regression with MCP regularization obtained by 5-fold cross-validation.

| Drug name | AUC | Sensitivity | Specivity | NPV | PPV |
|---|---|---|---|---|---|
| Kanamycin | 0,95 | 0,83 | 0,96 | 0,93 | 0,89 |
| Amikacin | 0,92 | 0,75 | 0,98 | 0,93 | 0,92 |
| Streptomycin | 0,95 | 0,84 | 0,94 | 0,91 | 0,88 |
| Ofloxacin | 0,90 | 0,70 | 0,95 | 0,91 | 0,81 |
| Moxifloxacin | 0,90 | 0,62 | 0,95 | 0,90 | 0,77 |
| Isoniazid | 0,98 | 0,90 | 0,99 | 0,96 | 0,97 |
| Rifampicin | 0,98 | 0,91 | 0,99 | 0,97 | 0,97 |
| Ethambutol | 0,95 | 0,75 | 0,96 | 0,96 | 0,75 |
| Pyrazinamide | 0,95 | 0,74 | 0,97 | 0,96 | 0,80 |
| Capreomycin | 0,88 | 0,65 | 0,96 | 0,91 | 0,80 |
| Ethionamide | 0,86 | 0,69 | 0,86 | 0,81 | 0,77 |
| Prothionamide | 0,72 | 0,58 | 0,74 | 0,72 | 0,61 |
| Ciprofloxacin | 0,98 | 0,90 | 0,99 | 0,97 | 0,96 |

Table S21. The quality metrics of logistic regression with L1 regularization obtained by 5-fold cross-validation.

| Drug name | AUC | Sensitivity | Specivity | NPV | PPV |
|---|---|---|---|---|---|
| Kanamycin | 0,96 | 0,83 | 0,95 | 0,93 | 0,88 |
| Amikacin | 0,92 | 0,75 | 0,98 | 0,93 | 0,92 |
| Streptomycin | 0,95 | 0,84 | 0,94 | 0,92 | 0,88 |
| Ofloxacin | 0,90 | 0,70 | 0,95 | 0,91 | 0,82 |
| Moxifloxacin | 0,90 | 0,62 | 0,95 | 0,90 | 0,78 |
| Isoniazid | 0,98 | 0,91 | 0,99 | 0,96 | 0,97 |
| Rifampicin | 0,98 | 0,91 | 0,99 | 0,97 | 0,97 |
| Ethambutol | 0,95 | 0,75 | 0,96 | 0,96 | 0,77 |
| Pyrazinamide | 0,95 | 0,73 | 0,97 | 0,96 | 0,79 |
| Capreomycin | 0,88 | 0,68 | 0,96 | 0,92 | 0,82 |
| Ethionamide | 0,86 | 0,68 | 0,86 | 0,80 | 0,76 |
| Prothionamide | 0,72 | 0,58 | 0,74 | 0,72 | 0,62 |
| Ciprofloxacin | 0,98 | 0,91 | 0,99 | 0,98 | 0,96 |

Table S22. The quality metrics of logistic regression with elastic net regularization obtained by 5-fold cross-validation.

| | AUC | Sensitivity | Specivity | NPV | PPV |
|---|---|---|---|---|---|
| Rifampicin | 0,98 | 0,89 | 0,99 | 0,96 | 0,96 |
| Isoniazid | 0,97 | 0,90 | 0,99 | 0,96 | 0,97 |
| Pyrazinamide | 0,95 | 0,73 | 0,97 | 0,96 | 0,79 |
| Ethambutol | 0,95 | 0,73 | 0,95 | 0,95 | 0,73 |
| Streptomycin | 0,95 | 0,83 | 0,94 | 0,91 | 0,88 |
| Kanamycin | 0,95 | 0,81 | 0,96 | 0,93 | 0,89 |
| Amikacin | 0,92 | 0,74 | 0,99 | 0,93 | 0,95 |
| Capreomycin | 0,87 | 0,65 | 0,97 | 0,91 | 0,84 |
| Ofloxacin | 0,90 | 0,69 | 0,96 | 0,91 | 0,86 |
| Moxifloxacin | 0,90 | 0,62 | 0,95 | 0,90 | 0,78 |
| Ciprofloxacin | 0,98 | 0,90 | 0,99 | 0,97 | 0,97 |
| Ethionamide | 0,87 | 0,65 | 0,86 | 0,79 | 0,76 |
| Prothionamide | 0,71 | 0,55 | 0,74 | 0,70 | 0,60 |

Table S23. The quality metrics of the dictionary method based on WHO catalog.

| Drug name | AUC | Sensitivity | Specivity | NPV | PPV |
|---|---|---|---|---|---|
| Amikacin | 0,86 | 0,74 | 0,98 | 0,93 | 0,91 |
| Capreomycin | 0,81 | 0,66 | 0,96 | 0,92 | 0,81 |
| Ethambutol | 0,89 | 0,84 | 0,93 | 0,97 | 0,69 |
| Ethionamide | 0,73 | 0,61 | 0,84 | 0,77 | 0,72 |
| Isoniazid | 0,93 | 0,88 | 0,99 | 0,95 | 0,98 |
| Kanamycin | 0,91 | 0,83 | 0,98 | 0,94 | 0,94 |
| Moxifloxacin | 0,81 | 0,69 | 0,94 | 0,92 | 0,76 |
| Pyrazinamide | 0,77 | 0,56 | 0,98 | 0,94 | 0,82 |
| Rifampicin | 0,95 | 0,93 | 0,98 | 0,98 | 0,95 |
| Streptomycin | 0,87 | 0,78 | 0,97 | 0,89 | 0,92 |
| Prothionamide | 0,60 | 0,24 | 0,95 | 0,76 | 0,64 |

Table S24. The comparison of stability of feature selection by different methods. The metrics are: the number of selected features averaged by dataset splits, the average fraction of majorly selected features from all features selected in each data split and the mean number of dataset splits in which the majorly selected features were selected. The maximal value of the metric for each drug is marked in bold.

[https://github.com/Reshetnikoff/m.tuberculosis-research-code/blob/main/Supplementary%20tables/Table%20S24.tsv](https://github.com/Reshetnikoff/m.tuberculosis-research-code/blob/main/Supplementary%20tables/Table%20S24.tsv)

Table S25. The clades selected by TreeBreaker for each drug and their composition. Each sample ID from the clades is presented in the last column with corresponding ID of the internal node from the second column which forms the clade. Some clades contain the other clades as subclades, so some sample IDs are repeated for the same drug.

[https://github.com/Reshetnikoff/m.tuberculosis-research-code/blob/main/Supplementary%20tables/Table%20S25.zip](https://github.com/Reshetnikoff/m.tuberculosis-research-code/blob/main/Supplementary%20tables/Table%20S25.zip)

Table S26. The descriptive statistics of TreeBreaker clades. For each drug the number of clades detected by TreeBreaker, the number of clades associated with location from the detected ones and the number of clades associated with lineages from the detected ones are presented.

| Drug | TreeBreaker clades | Associated with location | Associated with lineage |
|---|---|---|---|
| Ciprofloxacin | 18 | 0 | 8 |
| Prothionamide | 1 | 0 | 1 |
| Ethionamide | 12 | 1 | 10 |
| Kanamycin | 24 | 3 | 18 |

| | | | |
|---|---|---|---|
| Moxifloxacin | 10 | 3 | 6 |
| Amikacin | 26 | 10 | 21 |
| Capreomycin | 11 | 3 | 8 |
| Ofloxacin | 16 | 4 | 14 |
| Streptomycin | 47 | 18 | 36 |
| Pyrazinamide | 102 | 15 | 58 |
| Ethambutol | 92 | 20 | 64 |
| Isoniazid | 157 | 34 | 125 |
| Rifampicin | 163 | 28 | 111 |

Table S27. Locations and TBDB lineages of samples.

https://github.com/Reshetnikoff/m.tuberculosis-research-code/blob/main/Supplementary%20tables/Table%20S27.zip

Table S28. Locations and lineage characteristics of TreeBreaked clades. For each drug and each node_id of the clade root the number of resistant and susceptible isolates and the number of isolates for which the sampling location is known are presented. The last two columns present Bonferroni-corrected p-values of association of each clade with locations and lineages.

https://github.com/Reshetnikoff/m.tuberculosis-research-code/blob/main/Supplementary%20tables/Table%20S28.zip

Table S29. The features which were majorly selected by the ABESS algorithm trained using the feature set extended with features generated by TreeBreaker. All features were enumerated for each drug and the same IDs were assigned to the features constituting unified ones.

https://github.com/Reshetnikoff/m.tuberculosis-research-code/blob/main/Supplementary%20tables/Table%20S29.xlsx

Table S30. The features selected by the second iteration of ABESS. The features whose values fully coincide on the given dataset split were unified and were treated as one feature during the selection process. All features were enumerated for each drug and the same IDs were assigned to the features constituting unified ones.

https://github.com/Reshetnikoff/m.tuberculosis-research-code/blob/main/Supplementary%20tables/Table%20S30.zip

Table S31. The statistics of explained resistant isolates by the first and the second iteration of the HHS algorithm. We defined a resistant isolate to be explained by HHS on a given dataset split if logistic regression trained on the training partition of this split using the features majorly selected by HHS classifies this isolate as resistant. All the numbers were averaged by splits. In case of increase of the fraction of explained isolates for the given drug the values after the first and the second iteration are marked in bold.

| Drug name | Number of resistant isolates | Number of samples participated in the first ABESS iteration | Fraction of resistant samples explained by the first ABESS iteration | Number of resistant samples unexplained after the first ABESS iteration | Number of samples participated in the second ABESS iteration | Fraction of resistant samples explained by the first and the second ABESS iteration | Number of resistant samples unexplained after the second ABESS iteration |
|---|---|---|---|---|---|---|---|
| Rifampicin | 2277 | 9319 | **26,43%** | 1675 | 6856 | **88,19%** | 269 |
| Isoniazid | 2754 | 9252 | **17,68%** | 2267 | 7616 | **89,46%** | 290 |
| Pyrazinamide | 884 | 7334 | **33,26%** | 590 | 4894 | **81,95%** | 160 |
| Ethambutol | 1223 | 8143 | **4,86%** | 1164 | 7747 | **64,76%** | 431 |
| Streptomycin | 1344 | 3769 | **9,48%** | 1217 | 3411 | **25,64%** | 999 |
| Kanamycin | 275 | 961 | **81,98%** | 50 | 173 | **86,34%** | 38 |
| Amikacin | 322 | 1482 | **73,45%** | 86 | 393 | **74,69%** | 82 |
| Capreomycin | 323 | 1547 | **67,26%** | 106 | 506 | **69,06%** | 100 |
| Ofloxacin | 376 | 1559 | **56,38%** | 164 | 679 | **68,14%** | 120 |
| Moxifloxacin | 206 | 964 | **50,88%** | 101 | 473 | **53,31%** | 96 |
| Ciprofloxacin | 81 | 366 | **48,76%** | 41 | 187 | **65,35%** | 28 |
| Ethionamide | 222 | 553 | **63,54%** | 81 | 201 | **73,10%** | 60 |
| Prothionamide | 144 | 351 | **19,44%** | 116 | 282 | **36,81%** | 91 |

Table S32. The additional mutations majorly selected by HHS on the second iteration. The names of genes which were associated with resistance to given drug in previous studies (Table 2) assigned to the right drugs are marked in bold and underscored otherwise.

| Drug name | Gene | Position | Type of event | From | To | Feature ID | Number of splits in which the feature was selected |
|---|---|---|---|---|---|---|---|
| Rifampicin | ***rpoB*** | 435 | SNV | D | G | 0 | 5 |
| Rifampicin | ***rpoB*** | 445 | SNV | H | S | 1 | 4 |
| Rifampicin | ***rpoB*** | 432 | SNV | Q | K | 2 | 4 |
| Rifampicin | ***rpoB*** | 450 | SNV | S | L | 3 | 4 |
| Rifampicin | ***rpoB*** | 433 | Insertion | - | F | 4 | 4 |
| Rifampicin | *eccC2* | 905 | SNV | M | T | 5 | 4 |
| Rifampicin | *dnaJ2* | - | PF00226 is changed | - | - | 6 | 3 |
| Rifampicin | *nmtR* | -47 | SNV | T | C | 7 | 3 |
| Rifampicin | *mshD* | 71 | SNV | R | W | 8 | 3 |

| | | | | | | | |
|---|---|---|---|---|---|---|---|
| Rifampicin | *Rv3916c* | -62 | SNV | C | T | 9 | 3 |
| Rifampicin | ***rpoB*** | 445 | SNV | H | P | 10 | 3 |
| Rifampicin | *lppN* | -46 | SNV | A | G | 11 | 3 |
| Rifampicin | *gmhB* | - | Broken | - | - | 12 | 3 |
| Rifampicin | *suhB* | 263 | Deletion | VV | - | 13 | 3 |
| Isoniazid | ***katG*** | - | PF00141 is changed | - | - | 0 | 3 |
| Pyrazinamide | ***pncA*** | - | PF00857 is changed | - | - | 0 | 5 |
| Ethambutol | ***embB*** | 1002 | SNV | H | R | 0 | 5 |
| Ethambutol | ***embB*** | 306 | SNV | M | L | 1 | 5 |
| Ethambutol | ***embB*** | 306 | SNV | M | V | 2 | 5 |
| Ethambutol | ***embB*** | 306 | SNV | M | I | 3 | 3 |
| Streptomycin | ***rrs*** | 514 | SNV | T | G | 0 | 4 |
| Streptomycin | ***rpsL*** | 88 | SNV | K | R | 1 | 3 |
| Streptomycin | *Rv1465* | - | PF01592 is changed | - | - | 2 | 3 |
| Kanamycin | *rpoC* | - | PF00623 is changed | - | - | 0 | 3 |
| Amikacin | *Rv2168c* | -52 | Insertion | C | CAGCCGGGTCGTCACCGGCTGTCGGCGATAT | 0 | 3 |
| Amikacin | *mbtA* | - | PF00501 is changed | - | - | 1 | 3 |
| Capreomycin | *nanT* | - | PF07690 is changed | - | - | 0 | 3 |
| Capreomycin | *Rv2752c* | - | PF12706 is changed | - | - | 1 | 3 |
| Capreomycin | *Rv1986* | - | PF01810 is changed | - | - | 2 | 3 |
| Ofloxacin | *ald* | - | PF01262 is changed | - | - | 0 | 5 |
| Ofloxacin | *embA* | -16 | SNV | G | A | 1 | 5 |
| Ofloxacin | ***gyrB*** | - | PF01751 is changed | - | - | 2 | 4 |
| Ofloxacin | *hbhA* | 161 | Deletion | KKA | - | 3 | 3 |
| Moxifloxacin | *embA* | -12 | SNV | G | A | 0 | 4 |
| Moxifloxacin | *mdh* | - | PF02866 is changed | - | - | 1 | 3 |

| | | | | | | | |
|---|---|---|---|---|---|---|---|
| Moxifloxacin | *rpoC* | 527 | SNV | L | V | 2 | 3 |
| Moxifloxacin | *rocA* | - | PF00171 is changed | - | - | 3 | 3 |
| Ciprofloxacin | *rpoC* | - | PF00623 is changed | - | - | 0 | 3 |
| Ethionamide | ***ethA*** | - | PF00743 is changed | - | - | 0 | 4 |
| Ethionamide | *gnd1* | - | PF00393 is changed | - | - | 1 | 3 |
| Prothionamide | *nrdZ* | - | PF02867 is changed | - | - | 0 | 4 |
| Prothionamide | *Rv0566c* | - | PF04461 is changed | - | - | 1 | 3 |
| Prothionamide | *ndh* | - | PF07992 is changed | - | - | 2 | 3 |

Table S33. The features selected by the second iteration of HHS. All features were enumerated for each drug and the same IDs were assigned to the features constituting unified ones.

https://github.com/Reshetnikoff/m.tuberculosis-research-code/blob/main/Supplementary%20tables/Table%20S33.zip

### References


1. **Richardson A**. Logistic Regression: A Self-Learning Text, Third Edition by David G. Kleinbaum, Mitchel Klein. *Int Stat Rev* 2011;79:296–296.
2. **Tibshirani R**. Regression Shrinkage and Selection via the Lasso. *J R Stat Soc Ser B Methodol* 1996;58:267–288.
3. **Hoerl AE, Kennard RW**. Ridge Regression: Biased Estimation for Nonorthogonal Problems. *Technometrics* 1970;12:55–67.
4. **Hoerl AE, Kennard RW**. Ridge Regression: Applications to Nonorthogonal Problems. *Technometrics* 1970;12:69–82.
5. **Grogan TR, Elashoff DA**. A simulation based method for assessing the statistical significance of logistic regression models after common variable selection procedures. *Commun Stat Simul Comput* 2017;46:7180–7193.
6. **Saber MM, Shapiro BJ**. Benchmarking bacterial genome-wide association study methods using simulated genomes and phenotypes. *Microb Genomics* 2020;6:e000337.
7. **Lees JA, Mai TT, Galardini M, Wheeler NE, Horsfield ST, *et al.*** Improved Prediction of Bacterial Genotype-Phenotype Associations Using Interpretable Pangenome-Spanning Regressions. *mBio* 2020;11:e01344-20.
8. **Fan J, Li R**. Variable Selection via Nonconcave Penalized Likelihood and its Oracle Properties. *J Am Stat Assoc* 2001;96:1348–1360.
9. **Cun-Hui Zhang**. Nearly unbiased variable selection under minimax concave penalty. *Ann Stat* 2010;38:894–942.
10. **Kumar A, Bhattacharyya S, Bouchard K**. Numerical Characterization of Support

Recovery in Sparse Regression with Correlated Design. http://arXiv.org/abs/ (2021).
11. **Zhu J, Wen C, Zhu J, Zhang H, Wang X**. A polynomial algorithm for best-subset selection problem. *Proc Natl Acad Sci* 2020;117:33117.
12. **Libiseller-Egger J, Phelan J, Campino S, Mohareb F, Clark TG**. Robust detection of point mutations involved in multidrug-resistant Mycobacterium tuberculosis in the presence of co-occurrent resistance markers. *PLOS Comput Biol* 2020;16:e1008518.
13. **Zhang H, Li D, Zhao L, Fleming J, Lin N, *et al.*** Genome sequencing of 161 *Mycobacterium tuberculosis* isolates from China identifies genes and intergenic regions associated with drug resistance. *Nat Genet* 2013;45:1255–1260.
14. **Farhat MR, Shapiro BJ, Kieser KJ, Sultana R, Jacobson KR, *et al.*** Genomic analysis identifies targets of convergent positive selection in drug-resistant Mycobacterium tuberculosis. *Nat Genet* 2013;45:1183–1189.
15. **Coll F, Phelan J, Hill-Cawthorne GA, Nair MB, Mallard K, *et al.*** Genome-wide analysis of multi- and extensively drug-resistant Mycobacterium tuberculosis. *Nat Genet* 2018;50:307–316.
16. **Yang C, Luo T, Shen X, Wu J, Gan M, *et al.*** Transmission of multidrug-resistant Mycobacterium tuberculosis in Shanghai, China: a retrospective observational study using whole-genome sequencing and epidemiological investigation. *Lancet Infect Dis* 2017;17:275–284.
17. **Farhat MR, Freschi L, Calderon R, Ioerger T, Snyder M, *et al.*** GWAS for quantitative resistance phenotypes in Mycobacterium tuberculosis reveals resistance genes and regulatory regions. *Nat Commun* 2019;10:2128.
18. **Kouchaki S, Yang Y, Walker TM, Sarah Walker A, Wilson DJ, *et al.*** Application of machine learning techniques to tuberculosis drug resistance analysis. *Bioinformatics* 2019;35:2276–2282.
19. **Casali N, Nikolayevskyy V, Balabanova Y, Harris SR, Ignatyeva O, *et al.*** Evolution and transmission of drug-resistant tuberculosis in a Russian population. *Nat Genet* 2014;46:279–286.
20. **Pankhurst LJ, del Ojo Elias C, Votintseva AA, Walker TM, Cole K, *et al.*** Rapid, comprehensive, and affordable mycobacterial diagnosis with whole-genome sequencing: a prospective study. *Lancet Respir Med* 2016;4:49–58.
21. **Walker TM, Kohl TA, Omar SV, Hedge J, Del Ojo Elias C, *et al.*** Whole-genome sequencing for prediction of Mycobacterium tuberculosis drug susceptibility and resistance: a retrospective cohort study. *Lancet Infect Dis*. Epub ahead of print June 2015. DOI: 10.1016/S1473-3099(15)00062-6.
22. **Johnsen CH, Clausen PTLC, Aarestrup FM, Lund O**. Improved Resistance Prediction in Mycobacterium tuberculosis by Better Handling of Insertions and Deletions, Premature Stop Codons, and Filtering of Non-informative Sites. *Front Microbiol*;10. Epub ahead of print 2019. DOI: 10.3389/fmicb.2019.02464.
23. **Van der Auwera GA, Carneiro MO, Hartl C, Poplin R, del Angel G, *et al.*** From FastQ Data to High-Confidence Variant Calls: The Genome Analysis Toolkit Best Practices Pipeline. *Curr Protoc Bioinforma* 2013;43:11.10.1-11.10.33.
24. **Camacho C, Coulouris G, Avagyan V, Ma N, Papadopoulos J, *et al.*** BLAST+: architecture and applications. *BMC Bioinformatics* 2009;10:421.
25. **Cock PJA, Antao T, Chang JT, Chapman BA, Cox CJ, *et al.*** Biopython: freely available Python tools for computational molecular biology and bioinformatics. *Bioinformatics* 2009;25:1422–1423.
26. **Ge J, Li X, Jiang H, Liu H, Zhang T, *et al.*** Picasso: A Sparse Learning Library for High Dimensional Data Analysis in R and Python. *J Mach Learn Res* 2019;20:1–5.
27. **Fan J, Li R**. Variable Selection via Nonconcave Penalized Likelihood and its Oracle Properties. *J Am Stat Assoc* 2001;96:1348–1360.
28. **Zhang C-H**. Nearly unbiased variable selection under minimax concave penalty. *Ann Stat* 2010;38:894–942.
29. **Huerta-Cepas J, Serra F, Bork P**. ETE 3: Reconstruction, Analysis, and Visualization of Phylogenomic Data. *Mol Biol Evol* 2016;33:1635–1638.

30. **Phelan JE, O'Sullivan DM, Machado D, Ramos J, Oppong YEA, *et al.*** Integrating informatics tools and portable sequencing technology for rapid detection of resistance to anti-tuberculous drugs. *Genome Med* 2019;11:41.
