## Supplementary figures and images for "Feature selection and aggregation for antibiotic resistance GWAS in *Mycobacterium tuberculosis*: a comparative study"

### Fig. S1

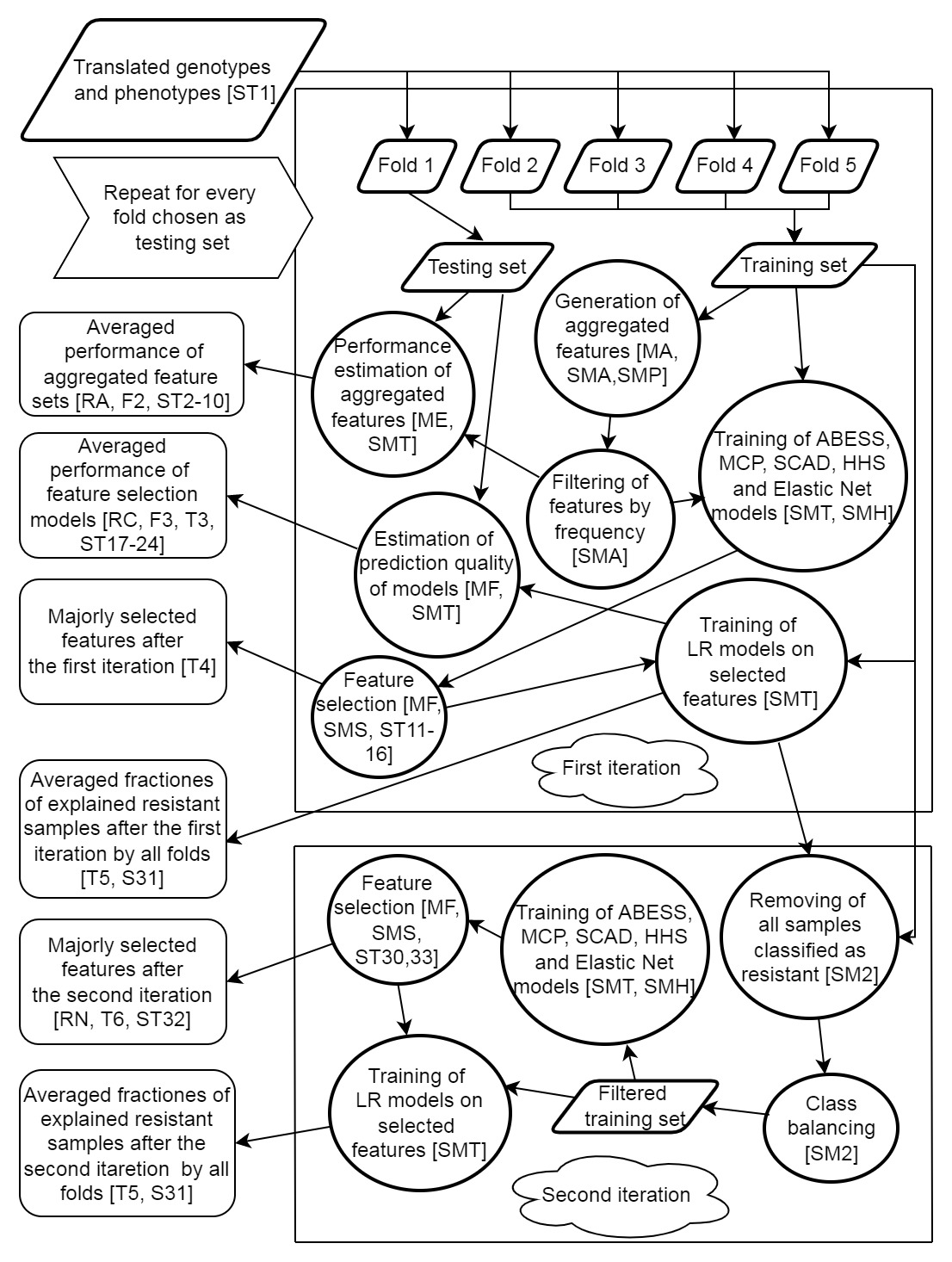

### Fig. S2

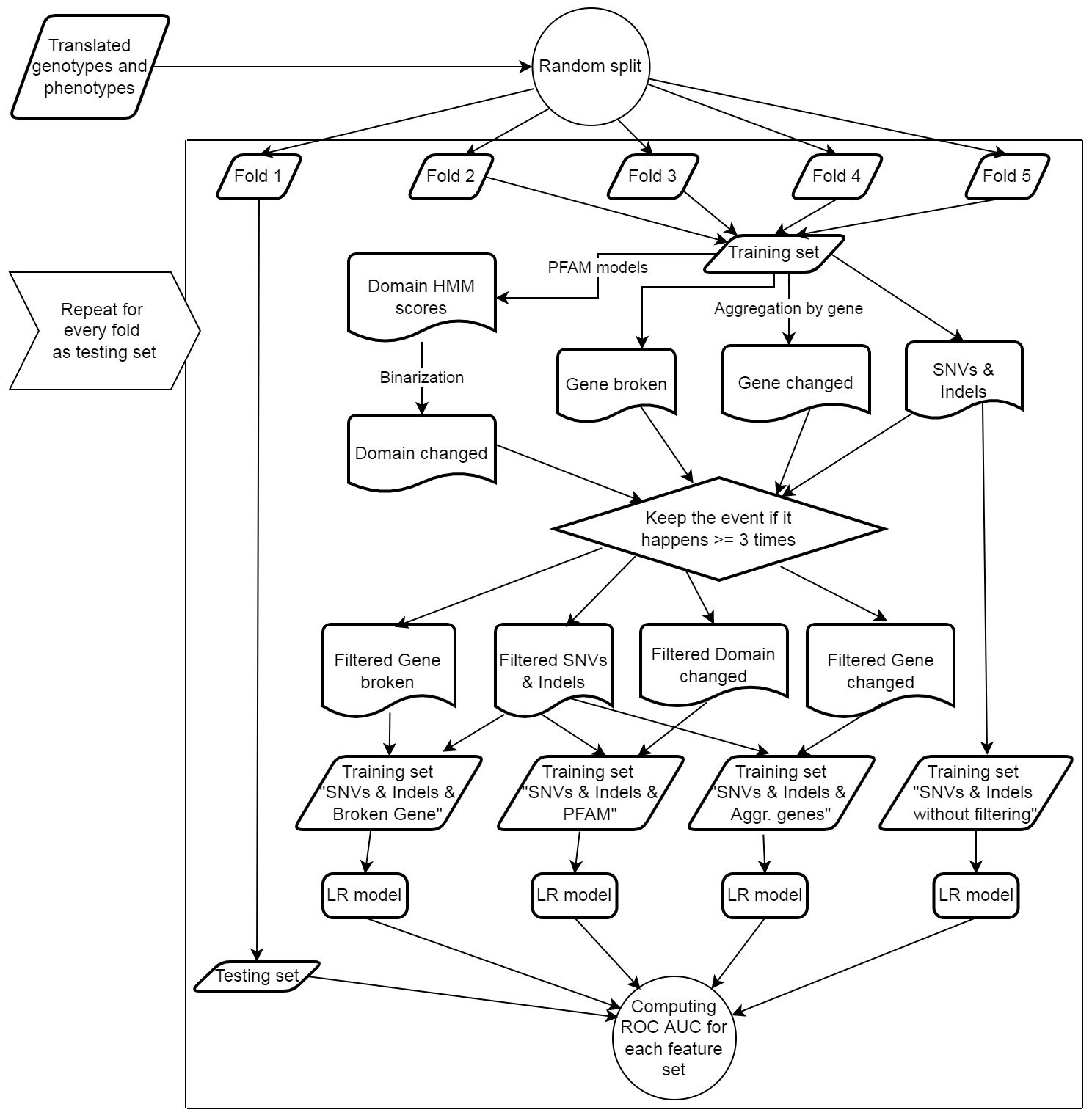
